# Using lineage-specific patterns to understand convergence of enzymatic functions led to the identification of Moraceae-specific P450s involved in furanocoumarin biosynthesis

**DOI:** 10.1101/2024.08.19.608558

**Authors:** Alexandre Bouillé, Rashmi Kumari, Alexandre Olry, Clément Charles, David R Nelson, Romain Larbat, Janet Thornton, Cloé Villard, Alain Hehn

## Abstract

- Specialized metabolites are molecules involved in plants interaction with their environment. Elucidating their biosynthetic pathways is a challenging but rewarding task, leading to societal applications and ecological insights. Furanocoumarins emerged multiple times in Angiosperms, raising the question of how different enzymes evolved into catalyzing identical reactions.
- To identify enzymes producing lineage-specific metabolites, an evolutionary-based approach was developed and applied to furanocoumarin biosynthesis in *Ficus carica* (Moraceae). This led to the characterization of CYP71B129-131a, three P450 enzymes whose evolution of the function was investigated using phylogenetics, structural comparisons and site-directed mutagenesis.
- CYP71B129 and CYP71B130,131a were found to hydroxylate umbelliferone (coumarin) and xanthotoxin (furanocoumarin), respectively. Results suggest that CYP71Bs xanthotoxin hydroxylase activity results from duplications and functional divergence of umbelliferone hydroxylase genes. Structural comparisons highlighted an amino acid affecting CYP71Bs substrate specificity, which may play a key role in allowing xanthotoxin hydroxylation in several P450 subfamilies.
- CYP71B130-131a characterization validates the proposed enzyme-discovery approach, which can be applied to different pathways and help to avoid the classic bottlenecks of specialized metabolism elucidation. The CYP71Bs also exemplify how furanocoumarin-biosynthetic enzymes can stem from coumarin-biosynthetic ones and provides insights into the molecular mechanisms underlying the multiple emergences of xanthotoxin hydroxylation in distant P450 subfamilies.

## Introduction

To adapt to their environment, plants have evolved various strategies including a large metabolic expansion which led to the production of myriads of molecules called plant specialized metabolites (PSMs) (Ono & Murata, 2023). Overall, the plant kingdom produces millions of structurally different PSMs, and each plant lineage possesses a unique metabolic profile. Remarkably, distant plants can produce similar PSMs, but rely on different enzymes to synthesize them. This phenomenon of convergent evolution towards an identical function is very common in specialized metabolism, making the biosynthesis of a given PSM potentially unique in each plant (Pichersky & Lewinsohn, 2011). Hence, PSM biosynthesis pathways are generally challenging to elucidate. To identify enzymes involved in PSM biosynthesis, various approaches were developed, relying on strategies such as sequence homology search, transcriptomic analyses, or gene mapping. For instance, it is common to analyze PSMs’ accumulation patterns (*e.g.*, tissues distribution, stress inducibility) to identify candidate enzymes displaying correlated gene expression (Ono & Murata, 2023). Such strategies are efficient and have already been successfully used, but they can be limited by factors such as low metabolite or transcription levels or recent divergent and convergent evolution (Pichersky & Lewinsohn, 2011). Promising advanced bioinformatic approaches such as deep learning and large-scale docking are also being developed, but still have limited efficiency (Daniel *et al*., 2015; Moore *et al*., 2019).

Furanocoumarins are a class of defensive PSM effective against various bioaggressors such as pathogens and phytophagous insects (Stevenson *et al*., 2003). Although derived from ubiquitous coumarins, furanocoumarins have only been identified in a few phylogenetically distant plant lineages including the Apiaceae, Rutaceae, Fabaceae and Moraceae (Sarker & Nahar, 2017). Over the past decades, the furanocoumarin biosynthesis pathway has been significantly elucidated in Apiaceous, Rutaceous and Moraceous species (**Fig. 1**). This pathway involves enzymes from the 2-oxoglutarate-dependent dioxygenase (Roselli *et al*., 2016), prenyltransferase (Munakata *et al*., 2020), O-methyltransferase (Hehmann *et al*., 2004), and cytochrome P450 (P450) families (Villard *et al*., 2021) (**Fig. 1**). Interestingly, identifying furanocoumarin-biosynthetic enzymes from different species showed that furanocoumarins are a case of convergent evolution: they emerged independently in distant plant lineages, in which they are synthesized by different enzymes (Limones-Mendez *et al*., 2020; Munakata *et al*., 2020; Villard *et al*., 2021).

**Figure 1:**
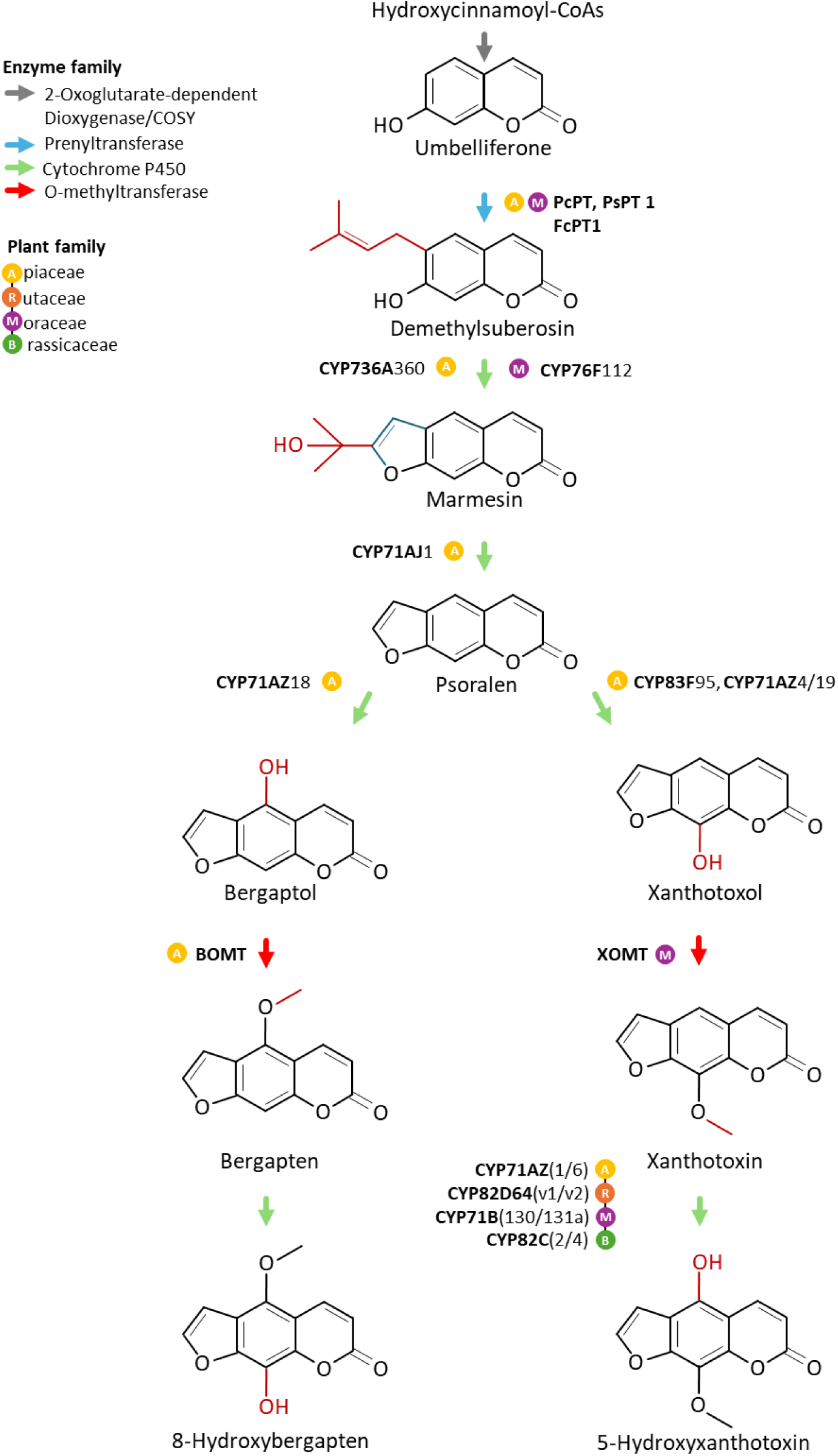
Simplified representation of the furanocoumarin biosynthetic pathway in Angiosperms. All furanocoumarin-biosynthetic enzymes described so far are detailed, together with associated plant families (Hehmann *et al*., 2004; Larbat *et al*., 2007; Kruse *et al*., 2008; Karamat *et al*., 2014; Roselli *et al*., 2016; Krieger *et al*., 2018; Limones-Mendez *et al*., 2020; Munakata *et al*., 2020; Villard *et al*., 2021; Wang *et al*., 2024; Ji *et al*., 2024). Abbreviations: PcPT, *Petroselinum crispum* prenyltransferase; PsPT1, *Pastinaca sativa* prenyltransferase 1; FcPT1, *Ficus carica* prenyltransferase 1; BOMT, bergaptol O-methyltransferase; XOMT, xanthotoxol O-methyltransferase.

Currently, the most referenced step of the furanocoumarin pathway corresponds to xanthotoxin hydroxylation (**Fig. 1**), catalyzed by 5-hydroxyxanthotoxin synthases (5OXHSs). All 5OHXSs characterized so far are P450s but belong to different (sub)families. Indeed, 5OHXSs from the Rutaceae (*i.e.*, *Citrus hystrix* and *Citrus paradisi*) are CYP82Ds (Limones-Mendez *et al*., 2020), while 5OHXS from the Apiaceae (*i.e.*, *Pastinaca sativa*) belong to the CYP71AZs (Krieger *et al*., 2018). Curiously, CYP82Cs from *Arabidopsis thaliana* (Brassicaceae) also display *in vivo* 5OHXS activity, although *Arabidopsis* does not naturally accumulate furanocoumarins (Kruse *et al*., 2008).

To pursue the elucidation of furanocoumarin biosynthesis and better understand the evolutionary process underlying the emergence of this pathway, we sought new furanocoumarin-biosynthetic enzymes in *Ficus carica* (Moraceae). To facilitate the identification of enzymes involved in the production of lineage-specific PSMs such as furanocoumarins, we established an evolutionary-based approach. Applying this approach, we compared the CYPomes of two Moraceae: *F. carica*, which produces furanocoumarins, and *Morus notabilis,* which as far as described does not. This led to identifying three *F. carica* CYP71Bs displaying 5OHXS and umbelliferone hydroxylase (UMBH) activity. Phylogenetics and comparative structural biology shed light on a lineage-specific diversification and highlighted an important residue which impacts enzyme activity and may have played a key role in the acquisition of the 5OHXS function.

## Materials and Methods

### Plant material and chemicals

*Ficus carica* plants were provided by INRAE Colmar (France). They were grown for four years at room temperature and under natural light. Standard specimens of phenolic compounds were purchased from various suppliers, as described in **Table S1**.

### Comparison of the CYP71 clan in *F. carica* and *M. notabilis*

A P450 nucleotide dataset was assembled with *F. carica* and *M. notabilis* CYP71 clan sequences (**Table S2**). *M. notabilis* sequences were retrieved from Ma *et. al.* (2014). *F. carica* sequences were identified by screening the PRJNA565858 reference genome (Usai *et al*., 2020) through BLASTN searches, using *M. notabilis* genes as queries and an e-value of 1000. Resulting hits were used to isolate full-length putative P450 genes and pseudogenes. Introns were predicted and removed using the NETGENE2 Server (http://www.cbs.dtu.dk/services/NetGene2/) (Hebsgaard, 1996). Putative pseudogenes containing frameshifts were altered by the addition of one or two “N” at the site of the indel(s) to recover a functional reading frame and allow subsequent analyses (**Table S2a**). *CYP51G* sequences were added to the dataset, to constitute the outgroup of the following tree. Sequences received a standardized name from the P450 nomenclature committee (https://drnelson.uthsc.edu/CytochromeP450.html, **Fig. S1**). Nucleotide sequences were then translated into proteins and aligned (**Fig. S2**) with MAAFT v.7 (https://mafft.cbrc.jp/alignment/server/) relying on default parameters (Kuraku *et al*., 2013; Katoh *et al*., 2019). Aligned sequences were used to build a Bayesian inference protein tree with MRBAYES v.3.2.7 (Ronquist & Huelsenbeck, 2003; Ronquist *et al*., 2012) at CIPRES v.3.3 (Cipres Science Gateway). The Bayesian Markov Chain Monte Carlo analysis was run with the following parameters: nst = mixed, rates = gamma, shape = (all), 2 million generations, two independent runs, four chains, temperature heating 0.05. The tree was drawn with FIGTREE v.1.4.4. Posterior probabilities supporting tree-branches were considered weak below 0.7, strong above 0.9.

### Cloning of *CYP71B129*, *CYP71B130* and *CYP71B131a* coding sequences

Total RNAs were extracted from young *F. carica* leaves and reverse-transcribed into complementary DNAs as described in Villard *et al*. (2021). *CYP71B129*, *CYP71B130* and *CYP71B131a* coding sequences (CDSs) were PCR-amplified using the PrimeSTAR® Max polymerase (Takara), according to supplier’s recommendations. Associated primers were 5’-GAAGAAAAATCATAGTAGAAATCTGAGTCATAAACTCCA-3’ (forward) and 5’-CAAATTTTTATTCTCTGTTTATATTTCTACTAGAAGATTGTTCAATGCAA-3’ (reverse) for *CYP71B129*, 5’-GGGAATCGAATTAGAAATCTAAGTGATAAGCTCCA-3’ (forward) and 5’-GTTTTTTCTAGAGAAAATTAACAACTTCGCTAACATATGTC-3’ (reverse) for *CYP71B130*, and 5’-GCGAATTGAACTAGAAATCAAACTGATAAGCTCCA-3’ (forward) and 5’-GTTACTAGTTTTATTTTTGAGAAAACATATTCAAGTCTATTTGAAGAAGTTT-3’ (reverse) for *CYP71B131a*. CDSs were then inserted into the pCR™8/GW/TOPO® TA vector (Invitrogen™, Thermo Fisher Scientific). Each gene was independently cloned and sequenced twice. A 3’-end 6xHis-tag was added by PCR, using the following primers: 5’-CCAATGGCTATTGATGCCCAAC-3’ (forward) and 5’CAACTAATGATGATGATGATGATGGAGGTGCTTCTTTGGGAC-3’ (reverse) for *CYP71B129*, 5’-CCAATGGCTATGGATGCCCAAC-3’ (forward) and 5’-GTTCTAATGATGATGATGATGATGTGCAAATAGGTGCTTCTTTG-3’ (reverse) for *CYP71B130*, and 5’-CCAATGGCTCTAGATGCCCG-3’ (forward) and 5’-GTTTTAATGATGATGATGATGATGTGCAAACAGGTGCTTCTTTG-3’ (reverse) for *CYP71B131a*. Finally, His-tagged CDSs were subcloned into the pYeDP60_GW® expression vector (Dueholm *et al*., 2015) using the Gateway LR Clonase™ II Enzyme Mix (Invitrogen™, Thermo Fisher Scientific).

### Synthesis of CYP71B mutants

Nucleotide sequences of His-tagged *CYP71B* mutants were chemically synthesized and subcloned into the pYeDP60 vector by GenScript Biotech (Netherlands). Cloning was performed using the Bam*HI* and Eco*RI* restriction enzymes.

### Heterologous expression of recombinant CYP71Bs

The pYeDP60 and pYedDP60-GW® vectors containing the various *CYP71Bs* genes, and the empty pYeDP60-GW® as control, were transformed into the WAT21 *Saccharomyces cerevisiae* strain according to Pompon *et al*. (1996) and Urban *et al*. (1997). P450 heterologous expression was performed as described in Larbat *et al*. (2007), with the following modifications: the yeasts were cultured at 18°C, and 30 mg. L^-1^ adenine was added in the YPGE medium (Kokina *et al*., 2014). Microsomal fractions containing heterologous P450s and P450 reductase *At*CPR2 were collected as described in Larbat *et al*. (2007). P450 expression was assessed by western blot, using 6xhistidine Epitope Tag antibodies (Acris, OriGene Technologies, USA, **Fig. S3**).

### *In vitro* characterization of CYP71Bs

A functional screening was performed on CYP71B129, CYP71B130, CYP71B131a and associated mutant enzymes. A total of 77 putative phenolic substrates were tested, including phenylpropenes, coumarins, furanocoumarins and pyranocoumarins (**Table S1**). For this, microsomal fractions were incubated at 28°C for 30 min in the presence of 200 µM substrate and 200 µM NADPH, in a final volume of 200 µL phosphate buffer, pH 7. Reactions were stopped by the addition of 65 µL acetonitrile/HCL 50:50 (1N). Reaction products were extracted twice with ethyl acetate, air dried, and resuspended in 100 µL methanol/water (80:20) for subsequent mass spectrometry analysis. Optimal pH and temperature were determined for CYP71B129, CYP71B130 and CYP71B131a; *Km_app_* were determined for all enzymes and mutants using CYP71B130 and CYP71B131a optimal conditions, as described in Villard *et al*. (2021).

### Identification and quantification of reaction products by mass spectrometry

Reaction products were analyzed using a UHPLC-MS system (Shimadzu, Japan), following the procedure described in Villard *et al*. (2021). Compounds were identified by comparing their retention time and *m/z* ratio to those of standard molecules, and quantified based on 320 nm UV signal, using the LABSOLUTION software (Shimadzu, Japan). Product identity was further confirmed using a UHPLC-HRMS system composed by a Thermo Vanquish liquid chromatographic system (Thermo Scientific) coupled to a tandem mass spectrometer (Orbitrap-IDX; Thermo Scientific). Analyses were performed as described in Villard *et al*. (2021), except for the gradient phase which was modified as follows (A/B; v/v): 10:90 between 0 and 2min, 30:70 at 10 min, 95:5 at 20 min, 95:5 between 20 and 25 min, and 10:90 from 26 to 30 min. Data were recorded and analyzed on the XCALIBUR^TM^ software (v.2.1.SP1. Build1160; Thermo Fisher Scientific).

### Phylogenetic analysis of Moraceous CYP71Bs

A CYP71B dataset (**Table S3a**) was constituted by performing a similarity search in the genomes and transcriptomic databases of 24 plant species from the Nitrogen Fixing Clade, available at GenBank (Benson *et al*., 2012). Genetic resources were screened through BLASTN search using *F. carica* CYP71B131a as query and an e-value of 1000. Full length putative P450 sequences were isolated, introns were removed, and putative pseudogenes were processed as explained earlier. Most sequences closer to *F. carica* CYP71AS33 than to *F. carica* CYP71B131a were removed. Putative genes received a standardized name from the P450 nomenclature committee (**Table S2a**). Previously characterized CYP71Bs (Schuhegger *et al*., 2006; Irmisch *et al*., 2014; Hansen *et al*., 2018) and a few representative CYP71ASs were added to the dataset. *F. carica* CYP71AH65 was added to constitute the outgroup of the following tree. Nucleotide sequences were aligned with GENEIOUS (Geneious prime 2019, **Fig. S4**), partitioned on MESQUITE v.3.6 (http://www.mesquiteproject.org) and used to build a Bayesian inference gene-family tree with MRBAYES v.3.2.7 (Ronquist & Huelsenbeck, 2003; Ronquist *et al*., 2012), as described in Villard *et al*. (2021). This first CYP71B gene-tree (**Fig. S5**) was used to reduce the initial CYP71B dataset to a smaller subset of Moraceae-specific sequences (**Table S3a**) that were realigned (**Fig. S5**) and used to generate a second tree, as described above. The trees were drawn with FIGTREE v.1.4.4. Posterior probabilities supporting tree-branches were considered weak below 0.7, strong above 0.9.

### Comparison of 5-hydroxyxanthotoxin synthases and closely related enzymes

A P450 dataset was assembled using the protein sequences of the eight 5OHXS enzymes described so far (*i.e.*, CYP71B130; CYP71B131a; CYP71AZ1 (GenBank EF127863.1); CYP71AZ6 (GenBank MH000221.1); CYP82C2 (Genbank NM_119348.2); CYP82C4 (GenBank: NM_119345.3); CYP82D64v1 (C. clementina v0.9 genome clementine0.9_007296m); CYP82D64v2 (C. sinensis DHSO v3.0 genome Cs_ont_1g009220.1) and seven closely related enzymes which do not display this activity (*i.e.*, CYP71B129; CYP71AZ3 (GenBank MH000218.1); CYP71AZ4 (GenBank MH000219.1); CYP71AZ5 (GenBank MH000220.1); CYP82C3, (GenBank NM_119346.3); CYP82D92 (Phytozome ID Ciclev10000846m); CYP82D179, (Phytozome ID Ciclev10000844m)). Protein sequences were aligned (**Fig. S6**) with MAAFT v.7 (https://mafft.cbrc.jp/alignment/server/) relying on default parameters (Kuraku *et al*., 2013; Katoh *et al*., 2019), and analyzed on MESQUITE v.3.6 (http://www.mesquiteproject.org).

### *In silico* modeling and docking experiments

CYP71B131a three-dimensional (3D) structures was folded using the ColabFold v1.5.2: AlphaFold2 (Mirdita *et al*., 2022), using mmseqs2_uniref_env and unpaired-paired MSA options, with standard options for monomeric structure predictions. To model the enzyme-substrate interactions, docking experiments were performed. First, a heme was docked into CYP71B131a 3D-structure. For this, CYP76AH1/miltiradiene (PDB ID: 7cb9) from *Salvia miltiorrhiza* (Gu *et al*., 2019), was used as a reference and overlapped with CYP71B131a 3D-structure to define a 18Å x 18Å x 18Å docking grid-box around the heme. Protein and heme files were prepared by adding polar hydrogens. A ‘+2’ charge was also added to the heme iron. Rigid docking of the heme (*i.e*., not allowing the protein to move) was performed using AutoDock Vina v1.2.3 (Trott & Olson, 2010). Subsequently, xanthotoxin and umbelliferone substrates were docked into the CYP71B131a-heme complex. The human microsomal P450 2A6 crystal structure (PDB ID: 1z11) was used as a reference (Yano *et al*., 2005), and overlapped with the CYP71B131a-heme complex to define a grid-box of 28Å x 24Å x 24Å around the xanthotoxin. Protein file was prepared by adding polar hydrogens, and a ‘+2’ charge was added to the heme iron. Residues within 5Å of xanthotoxin in the reference structure were identified as flexible docking residues. The ligand file was prepared by building bonds and adding polar hydrogen atoms. Flexible docking of the xanthotoxin or the umbelliferone was performed using AutoDock Vina v1.2.3 (Trott & Olson, 2010). CYP71B131a_G373A and CYP71B131a-G373S mutant enzymes were folded and substrates were docked as described above. Visualizations of the docking poses were prepared with PyMOL (DeLano, 2002).

## Results

### *Ficus*-*Morus* CYPome comparison highlights a CYP71B expansion

Since the furanocoumarin pathway appeared multiple times in higher plants, it is not advised to rely on classic similarity search to pursue its elucidation (Villard *et al*., 2021). Instead, to identify new enzymes involved in furanocoumarin biosynthesis, we developed a novel evolutionary-based approach. In previous work, we demonstrated that the marmesin synthase activity (**Fig. 1**) emerged relatively recently in the Moraceae, through a CYP76F expansion that occurred after the divergence of the *Ficus* and *Morus* ancestors (Villard *et al*., 2021). Therefore, we hypothesized that other *Ficus* enzymes catalyzing downstream furanocoumarin-biosynthetic reactions might have similar origins. This means other gene expansions that occurred after the *Ficus*-*Morus* divergence might enclose other furanocoumarin-biosynthetic enzymes. As target enzymes, we focused on P450s from the CYP71 clan, which play a central role in plant specialized metabolism (Nelson & Werck-Reichhart, 2011) and include all furanocoumarin-related P450s described so far (**Fig. 1**). Our approach therefore consisted in comparing the CYP71 clan in *F. carica* and *M. notabilis* to identify *Ficus*-specific expansions.

To identify *F. carica* CYP71 clan sequences, we screened the *F. carica* reference genome (Usai *et al*., 2020). A total of 133 full-length P450 sequences were isolated, corresponding to 85 genes and 48 pseudogenes (**Table S2a**). These sequences were classified into 18 families and 35 subfamilies of various sizes (**Table S2b**). Twelve families comprised a single subfamily or single *F. carica* sequence (*e.g.*, CYP84A). The six other families contained multiple subfamilies, with a maximum of 37 *F. carica* CYP71s distributed in eight subfamilies (**Table S2b**). Interestingly, most multi-sequence subfamilies result from tandem duplications and form physical gene clusters in *F. carica* genome. For instance, all nine *F. carica* CYP71Bs were clustered on a ≈115 kb fragment of chromosome 11 (**Table S2a**). *F. carica* P450s were then compared to the 97 *M. notabilis* CYP71 clan sequences, previously reported by Ma *et. al.* (2014). Even though more sequences were identified in *F. carica* (133) than *M. notabilis* (97), all 35 subfamilies observed in *F. carica* were present in *M. notabilis,* and six additional single-sequence subfamilies were only observed in *M. notabilis* (**Table S2b**).

*F. carica* and *M. notabilis* CYP71 clan sequences were then used to generate a protein-family phylogenetic tree (**Fig. 2**). Across this phylogeny, some families and subfamilies were highly conserved. For example, the CYP78A subfamily comprises six *F. carica* and *M. notabilis* pairs of orthologs, meaning that the last common *Ficus*-*Morus* ancestor already possessed six CYP78As. This analysis also revealed significant lineage-specific expansions of the CYP71B, CYP71AN, and CYP76F subfamilies in *F. carica*, and of the CYP82D subfamily in *M. notabilis* (**Fig. 2**). Since the CYP76F diversification gave rise to the marmesin synthase activity (Villard *et al*., 2021), we assumed the CYP71B and CYP71AN expansions might have given rise to other furanocoumarin-biosynthetic enzymes. Among these two subfamilies, the most drastic expansion occurred in the CYP71Bs, with one single *M. notabilis* sequence compared to nine *F. carica* orthologs (CYP71B129, CYP71B130, CYP71B131a, CYP71B131b, CYP71B133, CYP71B134, CYP71B135, CYP71B136, CYP71B137, **Fig. 2**). We therefore decided to focus on *F. carica* CYP71Bs for further characterization.

**Figure 2:**
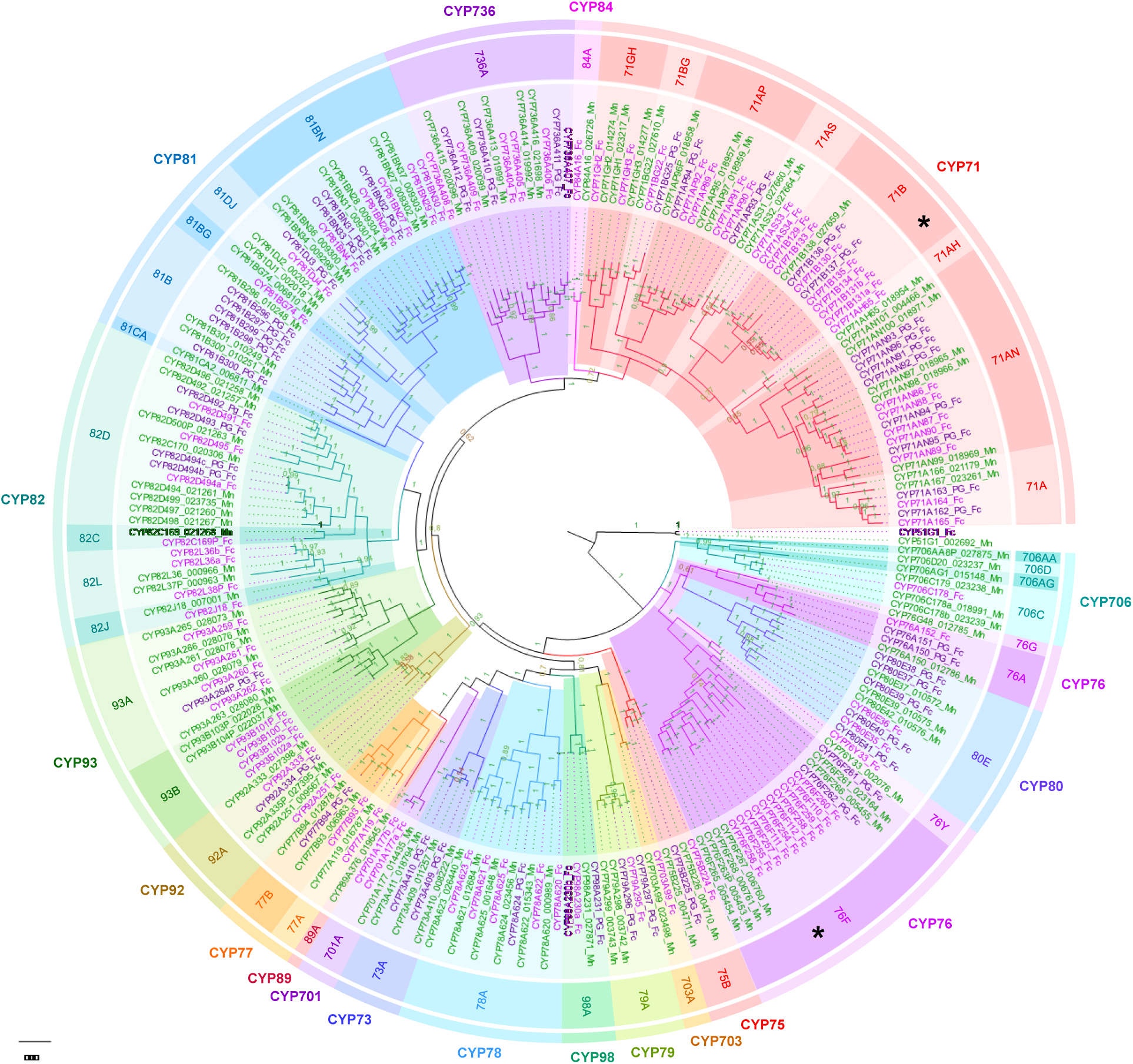
Comparison of the CYP71 clan in *F. carica* and M*. notabilis*. Bayesian inference tree of *F. carica* and *M. notabilis* CYP71 clan, rooted on the CYP51 clan. *F. carica* genes are colored in light purple, *F. carica* pseudogenes in dark purple, *M. notabilis* genes in green. For an easier understanding of the tree, clades are colored by P450 (sub)families, P450 (sub)family names are detailed, and the CYP76F and CYP71B subfamilies are highlighted with a black asterisk. *M. notabilis* sequence names include the accession numbers given in Ma *et al*. (2014).

### CYP71Bs include two 5-hydroxyxanthotoxin synthases and one umbelliferone hydroxylase

To investigate the function of the nine *F. carica* CYP71Bs, we cloned them from *F. carica* leaf RNA. Three genes were successfully amplified: *CYP71B129*, *CYP71B130* and *CYP71B131a*. CYP71B129-131a enzymes were heterologously produced in yeast (**Fig. S3**) and used to perform *in vitro* enzyme assays in the presence of NADPH and 77 putative substrates belonging to the coumarin, furanocoumarin, pyranocoumarin and phenylpropene families (**Table S1**). Results demonstrated that CYP71B130 and CYP71B131a were both able to convert xanthotoxin into 5-hydroxyxanthotoxin, making them 5OHXS enzymes (**Fig. 3; Fig. S7**). The identity of the product was formally identified by comparing its mass spectrometry characteristics to a commercial 5-hydroxyxanthotoxin standard.

**Figure 3:**
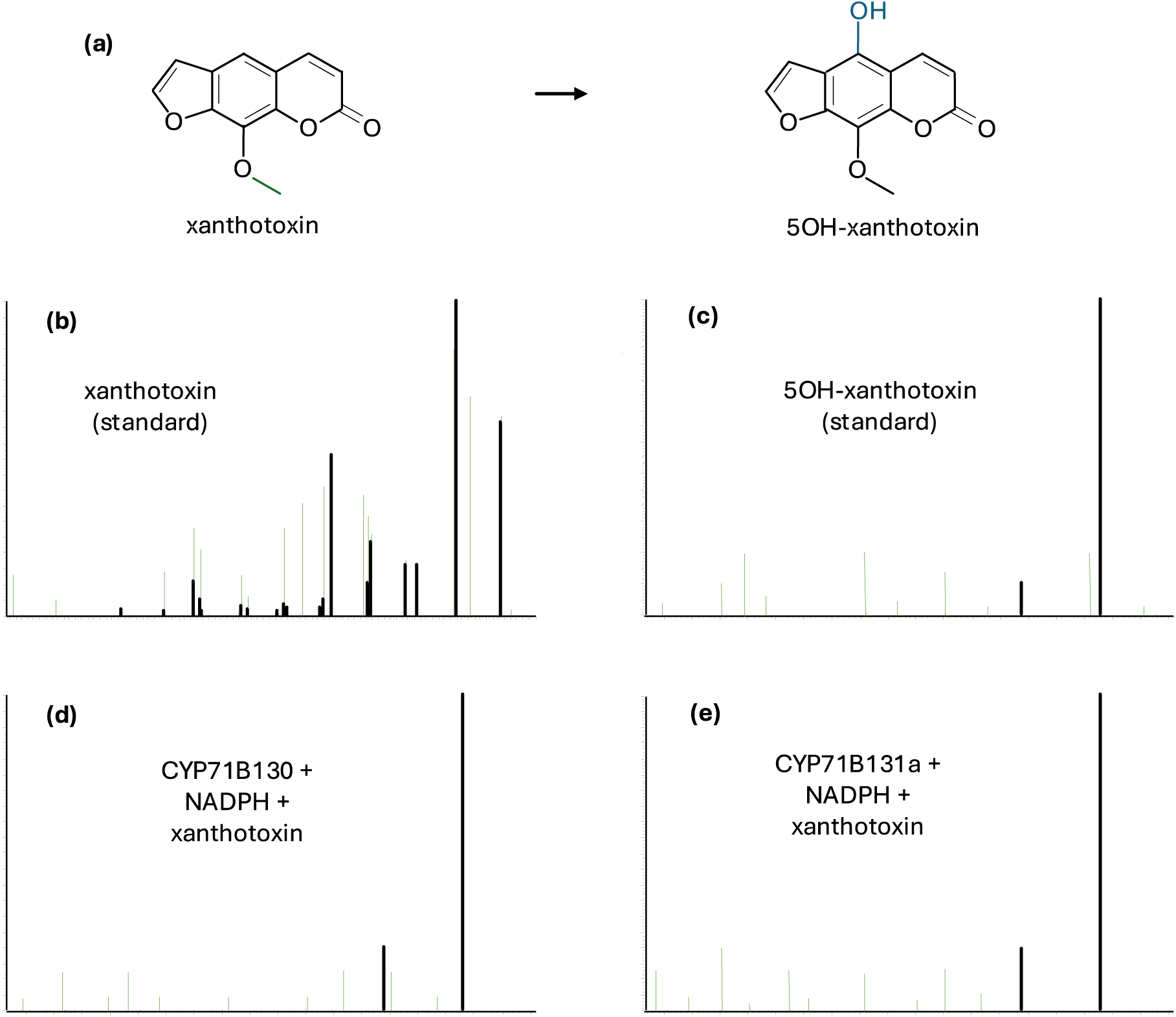
Conversion of xanthotoxin into 5-hydroxyxanthotoxin by CYP71B130 and CYP71B131a. (a) Molecular representation of the reaction. (b-e) Tandem mass spectrometry fragmentation patterns of the standards of (b) xanthotoxin and (c) 5-hydroxyxanthotoxin, and of the reaction products of (d) CYP71B130 and (e) CYP71B131a.

In contrast, CYP71B129 did not metabolize any of the tested furanocoumarins, but hydroxylated the coumarin umbelliferone (UMBH activity, **Fig. 4**). Theoretically, umbelliferone can be hydroxylated on five different carbons, leading to the production of 3,7-, 4,7-, 5,7-, 6,7- or 7,8-dihydroxycoumarin. Out of these five molecules, only four standards are commercially available, *i.e*., 4,7-, 5,7-, 6,7- and 7,8-dihydroxycoumarin. Using mass spectrometry (**Fig. S8**), we compared these four standards to CYP71B129 reaction product, showing that only 7,8-dihydroxycoumarin (daphnetin) had the same retention time as the product. This indicates that CYP71B129 produces either daphnetin or the untested 3,7-dihydroxycoumarin. As the amount of product synthesized by CYP71B129 was too low to consider an NMR investigation, we could not further confirm where the hydroxylation was performed.

**Figure 4:**
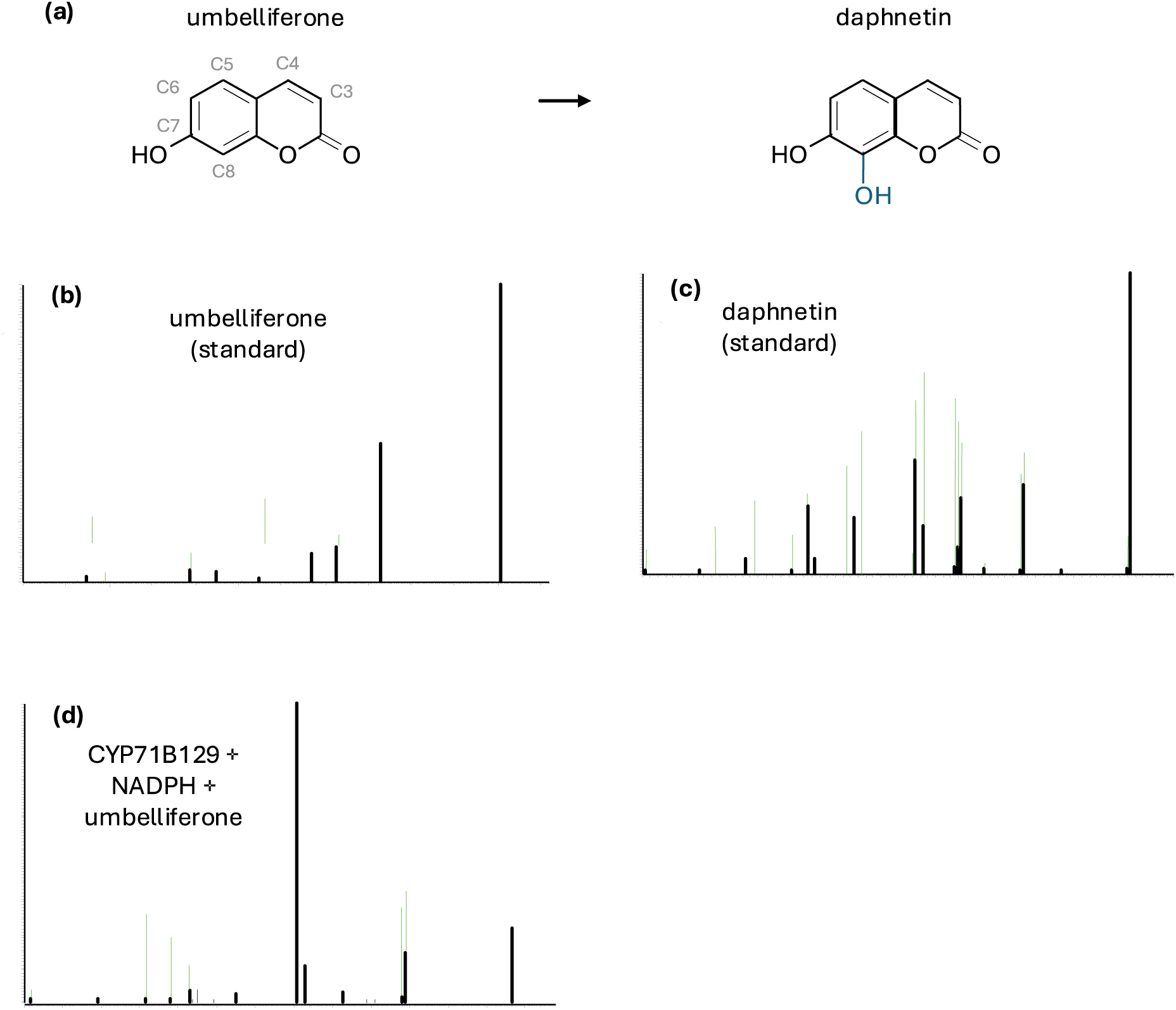
Metabolization of umbelliferone by CYP71B129. (a) Molecular representation of the reaction. Carbon numbering is indicated on the molecule. (b-d) Tandem mass spectrometry fragmentation patterns of the standards of (b) umbelliferone and (c) daphnetin, and (d) of the reaction product of CYP71B129.

Next, further enzymatic characterization showed that optimal activities of the three enzymes CYP71B129-131a with their respective substrates were obtained at pH 7.3-7.9 and a temperature of 24-26°C (**Fig. S10; Fig. S11**). The apparent *Km* values associated to these enzymes were also determined, showing that CYP71B130 (*Km*_app_ of 0.9 ± 0.7 µM) has a 10-times higher affinity for xanthotoxin than CYP71B131a (*Km*_app_ of 9.4 ± 1.2 µM) (**Fig. S12a-b**). The *Km*_app_ associated to umbelliferone hydroxylation by CYP71B129 was 28.5 ± 9.3 µM (**Fig. S12d**).

### The 5-hydroxyxanthotoxin synthase activity emerged in subclade B3 of Moraceous CYP71Bs

CYP71B129-131a are three *F. carica* enzymes sharing 83-88% nucleotide identity. However, CYP71B129 metabolizes a coumarin while CYP71B130 and CYP71B131a metabolize a furanocoumarin. To better understand the evolution of these enzymes, we performed a gene phylogenetic analysis. Screening 24 plant genomes, we identified 82 CYP71B sequences corresponding to 58 genes and 24 pseudogenes (**Table S3a**). Sequences were used to generate a gene-family phylogenetic tree (**Fig. S5**). On this gene-tree, all Rosales CYP71Bs form a well-supported clade which topology perfectly reflects the Rosales phylogeny (Zuntini *et al*., 2024). This indicates that the last common Rosales ancestor possessed a single CYP71B sequence, which either remained low-copy (*e.g.*, Rosaceae family) or diversified in a lineage-specific manner (*e.g.*, Moraceae family, **Fig. S5**).

Zooming on the gene-tree (**Fig. 5a**), we focused on Moraceae-specific CYP71Bs and classified them into two clades (A-B) comprising a total of four subclades (A, B1-B3). Clade A consists of all *Morus* and *Artocarpus* CYP71Bs. These CYP71Bs are well-conserved and present as single- or low-copy in the five screened *Morus* and *Artocarpus* genomes (**Fig. 5a**). Contrarily, clade B comprises all *Ficus* and *Antiaris* CYP71Bs. Each screened *Ficus* and *Antiaris* genome contained 4-14 CYP71B sequences, indicating significant expansion. Clade B is further subdivided into subclades B1-B3: each of these subclades contains both *Ficus* and *Antiaris* sequences, and therefore pre-dates the divergence of the *Ficus* and *Antiaris* ancestors (**Fig. 5a-b**). *F. carica* CYP71B129 belongs to subclade B2, CYP71B130 and CYP71B131a to subclade B3. The relatively low expansion and short branch-lengths observed in subclade B2 suggest that the CYP71B129 UMBH activity might not be a B2-specific novelty. Instead, it may be shared with clade A and subclade B1. Additional experiments are however required to support this hypothesis. Contrarily, sequences from subclade B3 diversified and accumulated more mutations, suggesting the 5OHXS activity observed in *F. carica* CYP71B130 and CYP71B131a emerged in this subclade. This means all B3-enzymes might be 5OHXSs.

**Figure 5:**
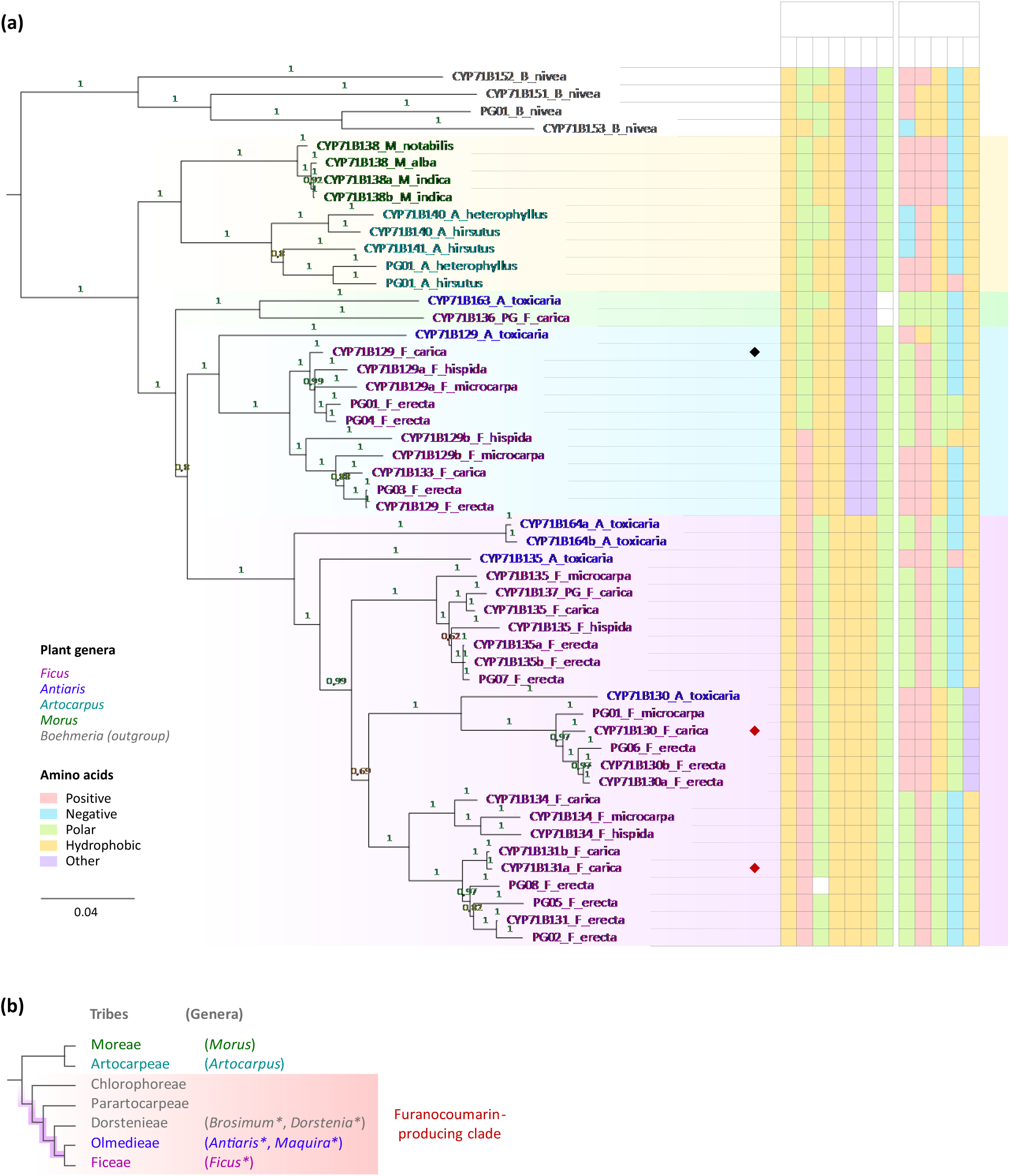
Emergence of the 5OHXS activity in the Moraceae family. (a) Gene-family tree of the CYP71Bs from the Moraceae family, rooted on CYP71Bs from *Boehmeria nivea* (Urticaceae). Characterized 5-hydroxyxanthotoxin synthase (5OHXS) are highlighted with a red diamond, characterized umbelliferone hydroxylase (UMBH) with a black diamond. Amino acids (AAs) of interest from the Substrate Recognition Sites (SRSs) are reported through the phylogeny, numbered according to *F. carica* CYP71B131a equivalent. Amino acids corresponding to CYP71B130 derived character states are highlighted in bold red, others are in gray. (b) Phylogeny of the Moraceae family, adapted from (Gardner *et al*., 2021). The tribes containing known furanocoumarin producing genera (*Ficus*, *Antiaris*, *Maquira*, *Brosimum* and *Dorstenia*) are highlighted in red (Rovinski *et al*., 1987; Vieira *et al*., 1999; Abegaz *et al*., 2004; Shi *et al*., 2014; Takahashi *et al*., 2017). The putative emergence of the 5OHXS activity in Moraceous CYP71Bs is highlighted in purple.

To further investigate the emergence of the 5OHXS activity within the CYP71Bs, we used our sequence alignment (**Fig. S7**) to compare the substrate recognition sites (SRSs, **Table S3b**) throughout the CYP71B phylogeny (Gotoh, 1992). Results highlighted seven SRS-residues (**Fig. 5a**) that were well-conserved in subclades A-, B1, and B2 (L116, Q211, A313, V318, P312, A373, T492, numbered according to CYP71B131a equivalents) but substituted in all B3-sequences (V116, H211, S313, I318, A372, G373, S492). These B3-specific synapomorphies might have played a key role in the emergence of the 5OHXS activity. Remarkably, SRS-residues were relatively well conserved throughout the B3 subclade, suggesting that even early branching B3-genes likely encode 5OHXSs (*i.e.*, *CYP71B135s*, *CYP71B164s*). For instance, only nine SRS-residues were different between the two characterized 5OHXS, *i.e*., H/Q104, K/R108, L/Q114, H/R248, Q/E249, I/V315, G/A320, L/I481 and A/T487 in CYP71B130/131a, respectively. These mutations probably explain the activity difference measured between the two enzymes. Among them, H104, K108, L115, Q248 and G320 are unique in subclade B3, as they are only observed in the six CYP71B130 sequences (**Fig. 5a**). These five recent mutations might thus be responsible for an increased 5OHXS activity in CYP71B130s.

Taken together, our results indicate that the 5OHXS activity emerged in subclade B3 of Moraceous CYP71Bs, probably from the UMBH activity. This evolution correlates with a few mutations affecting the SRSs. The 5OHXS activity therefore results from a lineage-specific CYP71B-diversification that happened after the divergence of the Ficeae and Moreae tribes, but before the divergence of the Ficeae and Olmedieae tribes. This pattern is consistent with the repartition of known furanocoumarin-producing species of the Moraceae family (**Fig. 5b**).

### 5-hydroxyxanthotoxin synthases possess a glycine in position 373

If the 5OHXS function emerged recently in the CYP71Bs, it has also been described in other P450 subfamilies, *i.e.*, CYP71AZ, CYP71B, CYP82C and CYP82D (Kruse *et al*., 2008; Krieger *et al*., 2018; Limones-Mendez *et al*., 2020). To understand 5OHXS independent emergence in distant P450 subfamilies, we compared the eight characterized 5OHXS enzymes (*i.e.*, CYP71AZ1, CYP71AZ6, CYP71B130, CYP71B131a, CYP82C2, CYP82C4, CYP82D64v1, CYP82D64v2) with seven closely related enzymes which do not exhibit this activity (*i.e.*, CYP71AZ3, CYP71AZ4, CYP71AZ5, CYP71B129, CYP82C3, CYP82D92, CYP82D179). For this, we aligned the protein sequences of the different enzymes and analyzed the SRSs throughout the alignment (**Fig. S7**). Overall, SRS-residues were either well-conserved across all sequences, varied according to P450 families or subfamilies, or varied without a clear pattern (**Table I**, **Fig. S7**). However, out of the 96 SRS-residues, residue 373 displayed a unique pattern (**Table I**, **Fig. S7**). Indeed, all investigated 5OHXS enzymes possessed a glycine in position 373 while non-5OHXS enzymes possessed either an alanine (CYP71B129, CYP71AZ3, CYP71AZ5, CYP82C3), a valine (CYP82D92) or a serine (CYP82D179). The only non-5OHXS enzyme possessing a glycine at this position was CYP71AZ4, which catalyzes another furanocoumarin-biosynthetic reaction (*i.e.*, xanthotoxol synthase) (Krieger *et al*., 2018). Additionally, G373 was also one of the seven B3-specific amino acids highlighted above (**Fig. 5a**). This unique and intriguing pattern hinted toward a critical role of residue 373 for the 5OHXS activity.

To explore this further, we docked the xanthotoxin substrate into CYP71B131a 3D-model (**Fig. 6a**) and defined the enzyme’s catalytic pocket as the 27 residues located at less than 5Å from the substrate (**Table 1**). These 27 residues form a subgroup of the 96 SRS-residues. Interestingly, residue 373 was included in this catalytic pocket, in very close proximity (≃3Å) to xanthotoxin’s furan ring (**Fig. 6d**).

**Figure 6:**
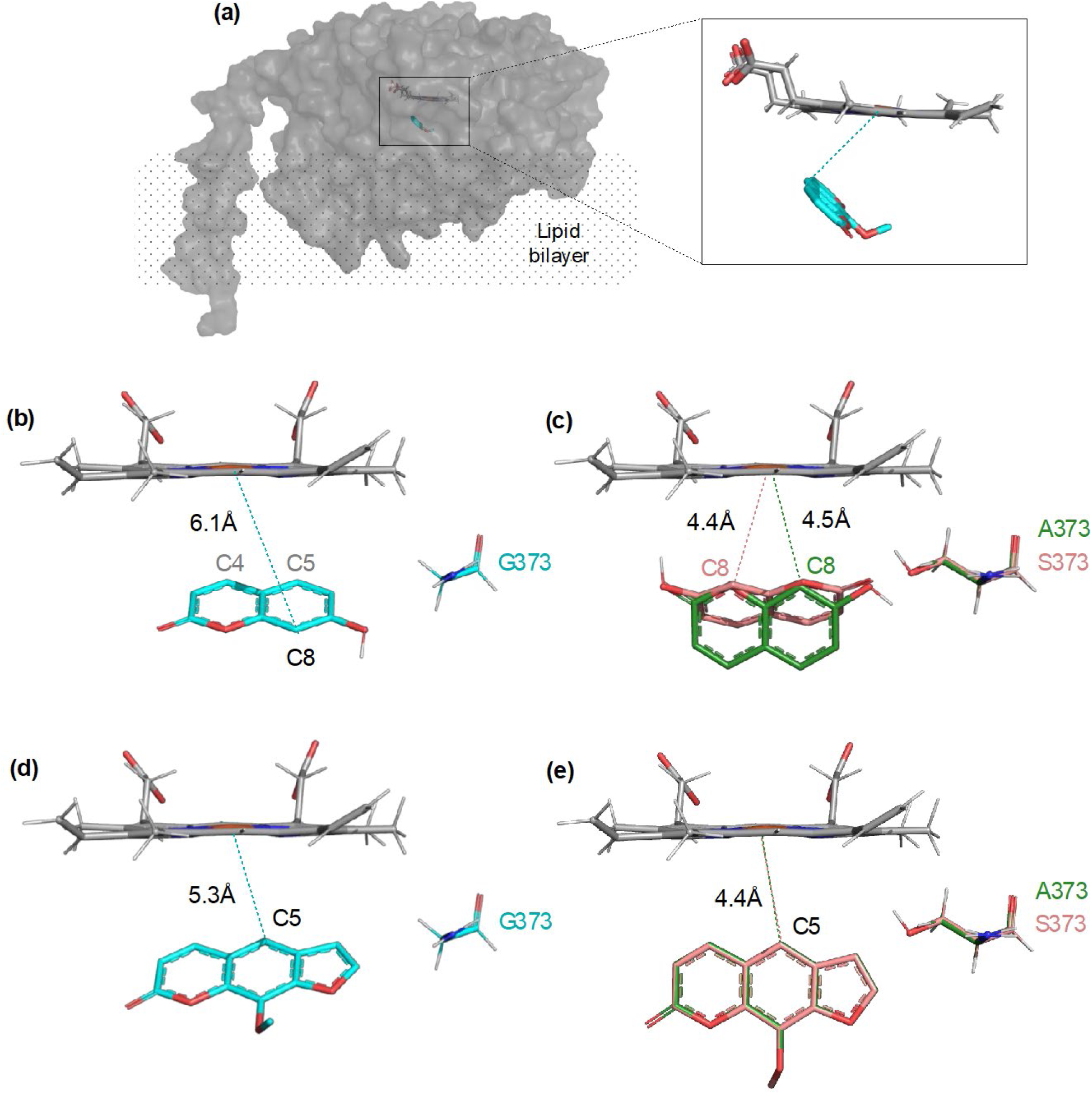
Docking of xanthotoxin and umbelliferone into CYP71B131a and associated mutant enzymes. (a) Three-dimensional (3D) model of CYP71B131a in complex with a heme (grey) and umbelliferone or xanthotoxin. (b-e) Most probable binding mode of umbelliferone (b,c) and xanthotoxin (d,e) into CYP71B131a (b,d) and associated mutants (c,e). Substrates are colored according to the enzyme they are docked in: blue for CYP71B131a, green for CYP71B131a_G373A, pink for CYP71B131a-G373S. Distance between substrates’ carbons and the heme iron are highlighted with dashed lines.

**Table 1:** Residues belonging to the substrate recognition sites (SRSs) and positioned close to the substrate (5Å) in 5-hydroxyxanthotoxin synthases (5OHXSs) and non-5OHXSs P450 enzymes. Residues conserved in most investigated sequences are highlighted in light blue, residues with a clear P450 family or subfamily patterns are highlighted in light yellow, residue varying according to P450 function is highlighted in light red, more variable residues are in white. Amino acids identical to the CYP71B131a equivalents are in bold red, others are in black. Residues are numbered according to the CYP71B131a equivalent.

| SRS |  | 1 |  |  |  |  |  |  |  |  |  |  | 2 |  | 3 | 4 |  |  |  |  |  |  | 5 |  |  |  |  | 6 |
| --- | --- | --- | --- | --- | --- | --- | --- | --- | --- | --- | --- | --- | --- | --- | --- | --- | --- | --- | --- | --- | --- | --- | --- | --- | --- | --- | --- | --- |
| Residues |  | 104 | 106 | 111 | 114 | 115 | 116 | 117 | 118 | 120 | 121 | 122 | 213 | 217 | 247 | 303 | 304 | 305 | 307 | 308 | 309 | 312 | 372 | 373 | 376 | 377 | 379 | 491 |
| 5OHXS | CYP71B130 | H | A | Y | L | D | V | A | F | P | Y | G | A | M | F | L | N | M | L | G | G | T | A | G | L | L | R | L |
|  | CYP71B131a | Q | A | Y | Q | D | V | A | F | P | Y | G | A | M | F | L | N | M | L | G | G | T | A | G | L | L | R | L |
|  | CYP71AZ1 | T | M | Y | L | D | V | A | F | P | Y | S | L | L | F | L | N | V | N | G | G | T | T | G | L | I | R | I |
|  | CYP71AZ6 | T | M | Y | L | D | V | A | F | T | Y | S | I | L | F | L | N | V | N | G | G | T | T | G | L | I | R | L |
|  | CYP82D64v1 | V | G | Y | S | L | L | G | F | P | Y | G | F | M | A | L | A | L | L | G | G | T | A | G | L | A | R | L |
|  | CYP82D64v2 | V | G | Y | S | L | L | G | F | P | Y | G | F | M | A | L | A | L | L | G | G | T | A | G | L | A | R | L |
|  | CYP82C2 | A | A | Y | A | V | F | G | F | P | Y | S | F | V | G | L | A | L | L | G | G | T | A | G | L | G | R | L |
|  | CYP82C4 | A | A | Y | A | V | F | G | F | P | Y | S | F | I | G | L | A | L | L | G | G | T | A | G | L | G | R | L |
| Non-5OHXS | CYP71B129 | Q | A | Y | Q | D | L | A | F | P | Y | G | A | M | F | L | N | M | L | G | G | T | P | A | L | L | R | L |
|  | CYP71AZ3 | T | M | Y | L | D | V | A | F | P | Y | T | I | F | F | L | N | V | N | G | G | T | P | A | L | V | R | L |
|  | CYP71AZ4 | T | S | Y | L | E | M | A | F | P | Y | G | I | F | F | L | N | V | S | G | G | T | P | G | L | I | R | L |
|  | CYP71AZ5 | T | T | Y | L | D | M | A | F | P | Y | S | L | S | F | Q | N | I | N | G | G | T | P | A | L | L | R | - |
|  | CYP82D92 | L | M | Y | S | M | F | G | F | P | Y | G | S | S | A | L | G | L | L | A | A | T | A | V | L | V | H | L |
|  | CYP82D179 | L | M | Y | S | M | F | G | F | P | Y | G | F | S | A | M | G | L | L | A | A | N | A | S | S | I | H | L |
|  | CYP82C3 | A | A | Y | - | - | - | - | - | - | - | - | F | V | R | L | A | L | L | G | G | T | A | A | L | G | R | L |

### Residue 373 strongly impacts CYP71B131a enzymatic activity

To experimentally investigate the impact of amino acid 373 on the 5OHXS enzyme activity, we designed a series of CYP71B131a mutants, in which G373 was substituted with an alanine (A), serine (S) or valine (V). These substitutions correspond to the amino acids observed in the non-5OHXS P450 enzymes described above. Each substitution introduces a gradual steric hindrance (G<A<S<V) and/or variable polarity (S). CYP71B131a mutant genes were synthesized, and associated enzymes were expressed and characterized. Results showed that CYP71B131a_G373A and CYP71B131a_G373S could still catalyze the 5OHXS reaction (**Fig. S7**). The affinity of CYP71B131a_G373A for xanthotoxin was 3-times lower than that of the wild-type enzyme (*Km_app_* = 20.8 ± 8µM). The activity of CYP71B131a_G373S was too low to accurately determine its *Km_app_*. For CYP71B131a_G373V, no 5OHXS activity was detected. The three mutant enzymes were also incubated in the presence of umbelliferone. Interestingly, they all exhibited the UMBH activity (**Fig. S8**), and showed a rather high affinity for umbelliferone (*Km_app_* = 9.3 ± 8.9 µM for CYP71B131a_G373A; 12.3 ± 9.9 µM for CYP71B131a-G373S, **Fig. S11e-f**. Next, to test if a glycine in position 373 would be enough to recover the 5OHXS activity, we designed a CYP71B129_A373G mutant and characterized the associated enzyme. No 5OHXS activity was detected, and CYP71B129_A373G still exhibited traces of UMBH activity (**Fig. S8h**).

To understand the impact of the G373A and G373S mutations on CYP71B131a 5OHXS activity, we built 3D-models of the associated mutant enzymes and performed docking experiments with xanthotoxin and umbelliferone. Xanthotoxin dockings showed that substituting G373 by an alanine or a serine slightly pushes the xanthotoxin and modifies its orientation toward the heme (**Fig. 6e**). Given the experimental results, this altered position might not be favorable for xanthotoxin binding. In the case of umbelliferone, amino acid 373 had an even stronger impact. Indeed, the best docking state obtained with CYP71B131a positioned umbelliferone with its carbons C4 and C5 oriented toward the heme iron. In contrast, docking performed with CYP71B131a_G373A and CYP71B131a_G373S resulted in a horizontal rotation of umbelliferone, positioning carbon C8 towards the heme iron (**Fig. 6c**). These results strongly suggest that CYP71Bs’ UMBH activity corresponds to a C8 hydroxylation, and thus, to daphnetin production. This would also explain why the wild-type CYP71B131a does not exhibit this activity.

In summary, our results demonstrated that amino acid 373 plays an important role in shaping the CYP71B131a active site and directly impacts substrate specificity and affinity. Indeed, increased steric hindrance at position 373 influences the positioning of the substrate in the active site, resulting in a decreased affinity for xanthotoxin and an increased affinity for umbelliferone. However important, amino acid 373 was not critical for the acquisition of the 5OHXS function in CYP71Bs, as CYP71B131a_G373A still exhibited 5OHXS activity while CYP71B129_A373G did not. This suggests additional residues must also play an essential role in the 5OHXS activity.

## Discussion

PSMs are molecules of ecological and societal interest with applications in medicine, agronomy, cosmetics, and in the agri-food industry (Weng *et al*., 2021). As plants do not usually produce enough PSMs quantities to cover large-scale applications, a major challenge of the field consists in elucidating PSM biosynthetic pathways, to help to develop biotechnological tools for large-scale production (Ono & Murata, 2023). This also provides new insights into plant biochemistry and can help better understand PSM ecological roles.

In this study, we developed an evolutionary-based approach to identify enzymes involved in the biosynthesis of lineage-specific PSMs, and we applied it to furanocoumarin biosynthesis. For this, we compared the genes of two Moraceae with different PSM profiles: *F. carica* and *M. notabilis*. As far as we know, furanocoumarins are indeed present in *Ficus* but not in *Morus*, and likely appeared in Moraceae after the *Ficus-Morus* divergence (Villard *et al*., 2021). To identify furanocoumarin biosynthetic specific genes, we thus looked for lineage-specific gene expansions which could correlate with furanocoumarin lineage-specific emergence. This led to the identification of a CYP71B diversification, amongst which two enzymes exhibited the 5OHXS activity. Our approach therefore proved successful and could be generalized to other families of PSMs, enzymes, and plants, provided that the target PSM is lineage-specific and its repartition in plants is known. If phylogenetics is widely used to analyze characterized-enzymes, it has so far rarely been used to select candidate-enzymes, and only as a minor tool complementing classic approaches (Pluskal *et al*., 2019; Hodgson *et al*., 2019). Our proposed approach is therefore original but supported by many examples of lineage-specific enzymes producing lineage-specific PSMs, as reviewed by Pichersky and Lewinsohn (2011). Moreover, as it only requires genomic resources which are now plentiful and easy to generate, it can be applied to virtually any lineage-specific PSM, regardless of previously limiting factors such as convergent evolution and low PSM production. Our approach therefore has great potential to help address the classic bottlenecks of specialized enzyme discovery. Of course, it also has limits: for instance, it cannot highlight newly functionalized enzymes that evolved without prior major gene duplications. However evolving new functions without gene duplications is relatively rare (Yu *et al*., 2015). Our approach thus constitutes an excellent tool to elucidate PSM biosynthesis pathways and can be used in combination with other existing approaches. Finally, it also means that the CYP71AN expansion observed in *F. carica* likely includes enzymes involved in the production of furanocoumarins or another lineage-specific PSM.

Experimental results showed that CYP71B129-131a are highly selective enzymes hydroxylating umbelliferone (CYP71B129) or converting xanthotoxin into 5-hydroxyxanthotoxin (CYP71B130,131a). The affinity of CYP71B130 (*Km_app_* = 0.9 ± 0.7 µM) and CYP71B131a (*Km_app_* = 9.4 ± 1.2 µM) for xanthotoxin falls within the classic affinity range determined for other furanocoumarin-biosynthetic P450s (*i.e.*, *Km_app_* ≈ 0.5-13 µM for CY71AJ1-4, CYP71AZ1,4,6, CYP82D64) (Krieger *et al*., 2018; Limones-Mendez *et al*., 2020). The only exception so far is CYP76F112, which displays a very high affinity (*Km_app_* = 32.2 ± 3.9 nM) for demethylsuberosin (Villard *et al*., 2021). On the contrary, P450s involved in coumarin biosynthesis often have lower affinity for their substrate. This is for instance, the case of CYP71AZ3, which displays a rather low affinity (*Km_app_* = 248.6 ± 51.9 μM) for esculetin (Krieger *et al*., 2018). This is also consistent with CYP71B129 slightly lower affinity for umbelliferone (*Km_app_* = 28.5 ± 9.3 µM). In addition, according to the Kitajima *et al. F. carica* RNAseq libraries, *CYP71B130* and *CYP71B131a* are overexpressed in petiole latex (Kitajima *et al*., 2018), which has a high furanocoumarin content (Munakata *et al*., 2020). Taken together, the high selectivity and relatively good affinity of CYP71B129-131a, their expression patterns, and the presence of both umbelliferone and xanthotoxin substrates in *Ficus carica* (Mamoucha *et al*., 2016), suggest that the UMBH and 5OHXS activities might be CYP71B129 and CYP71B130-131a physiological functions, respectively. However, to the best of our knowledge, associated products were never reported in *Ficus*. This could be because these products are not often included in metabolomic studies, or because they do not accumulate in *Ficus* but are instead quickly converted into different compounds.

Furanocoumarins are lineage-specific PSMs which derive from ubiquitous coumarins such as umbelliferone (**Fig. 1**) (Munakata *et al*., 2020). Interestingly, several furanocoumarin-biosynthetic enzymes were found to metabolize coumarins and/or to be closely related to coumarin-biosynthetic enzymes(Kruse *et al*., 2008; Rajniak *et al*., 2018; Krieger *et al*., 2018; Limones-Mendez *et al*., 2020).. For instance, within the CYP71AZ subfamily, CYP71AZ3 metabolizes esculetin (coumarin), CYP71AZ4 metabolizes scopoletin (coumarin) and psoralen (furanocoumarin), while CYP71AZ1 and CYP71AZ6 metabolize xanthotoxin (furanocoumarin) (Krieger *et al*., 2018). In this study, we observed a similar pattern in the CYP71B subfamily, as it includes both UMBH (coumarin) and 5OHXS (furanocoumarin) activities. Moreover, our phylogenetic analyses strongly suggest that CYP71B 5OHXS activity evolved from the UMBH activity, via duplications and functional divergence of the *UMBH* genes. As closely related P450s often metabolize similar compounds (Nelson & Werck-Reichhart, 2011), CYP71B functional divergence was likely facilitated by the structural similarity between umbelliferone and xanthotoxin. The CYP71B subfamily therefore perfectly exemplifies how furanocoumarin-biosynthetic enzymes can stem from coumarin-biosynthetic ones.

In previous studies, CYP71AZ1, CYP71AZ6, CYP82C2, CYP82C4, CYP82D64v1 and CYP82D64v2 were described as 5OHXS enzymes (Kruse *et al*., 2008; Krieger *et al*., 2018; Limones-Mendez *et al*., 2020). With the characterization of CYP71B130 and CYP71B131a, the 5OHXS activity has now been identified in four P450 subfamilies. To better understand how multiple P450 subfamilies acquired the ability to metabolize xanthotoxin, we decided to perform SRS analyses. The SRSs are small regions of P450 enzymes known to regulate substrate specificity. Hence, amino acid substitutions affecting the SRSs often result in significant activity changes such as substrate range, affinity, regio- and stereoselectivity (Gotoh, 1992; Villard *et al*., 2021). First, we focused on the CYP71B subfamily and identified seven SRS-amino acids that were well-conserved in most Moraceous CYP71Bs but substituted in all putative 5OHXSs from subclade B3 (*i.e.*, substitutions L116V, Q211H, A313S, V318I, P372A, A373G, T492S). In CYP76F112, similar evolutionary patterns were described for four SRS-amino acids that were experimentally shown to impact substrate specificity and affinity (Villard *et al*., 2021). It can thus reasonably be assumed that (some of) the seven 5OHXS-specific SRS-substitutions identified in the CYP71Bs played a key role in the substrate specificity switch from umbelliferone to xanthotoxin. Then, we extended our analysis to the CYP71B, CYP71AZ, CYP82C and CYP82D subfamilies and identified an SRS-amino acids displaying a unique pattern: all 5OHXSs from the four subfamilies possessed a glycine in position 373, while non-5OHXS enzymes possessed a different amino acid at this position. Experimental results showed that increased steric hindrance at position 373 lowered CYP71B131a affinity for xanthotoxin and allowed umbelliferone metabolization. These results are consistent with the work of Krieger *et al*. (2018), in which the simultaneous substitution of seven SRS-amino acids, including position 373, impacted CYP71AZ4 substrate specificity. This suggests residue 373 may contribute to substrate specificity in multiple P450 subfamilies. In the CYP71B subfamily, UMBH enzymes possess an alanine in position 373, while 5OHXS possess a glycine (**Fig. 5a**). However, substituting CYP71B129 A373 by a glycine did not allow xanthotoxin metabolization, while substituting CYP71B131a G373 by an alanine was not enough to suppress the 5OHXS activity. Taken together, our results allow us to conclude that the acquisition of the 5OHXS activity in the CYP71Bs, CYP71AZs, CYP82Cs and CYP82Ds was made possible by the substitution of different sets of amino acids in each subfamily, tailored per enzymatic environment. Amongst these substitutions, the only constant through the four subfamilies is G373. This means that each enzyme subfamily took a different evolutive path leading to the 5OHXS activity, but along each path, G373 was found to be favorable to xanthotoxin metabolization. Although not indispensable, G373 has thus played a key role in the independent emergence of the 5OHXS activity in different P450 subfamilies.

From a different perspective, phylogenetic analyses indicated that the 5OHXS activity emerged recently in the Moraceae family, through a lineage-specific CYP71Bs diversification. This evolutionary pattern is similar to that of CYP76F112, the *F. carica* marmesin synthase (Villard *et al*., 2021). When CYP76F112 was identified, the only Moraceous genomes available were from *Ficus* and *Morus*. By including recent *Artocarpus* and *Antiaris* genomes, we could strengthen and complete previous hypotheses. Indeed, the Moraceae family is divided into two clades (**Fig. 5b**): the first one consists of the Moreae (*e.g.*, *Morus*) and Artocarpeae (*e.g.*, *Artocarpus*) tribes (Gardner *et al*., 2021), in which furanocoumarins were never reported. The second clade comprises five tribes, including the Dorstenieae (*e.g.*, *Dorstenia*), Olmedieae (*e.g.*, *Antiaris*) and Ficeae (*e.g.*, *Ficus*), in which furanocoumarins were identified (Gottlieb *et al*., 1972; Rovinski *et al*., 1987; Heinke *et al*., 2011; Shi *et al*., 2014; Takahashi *et al*., 2017). This clade will be defined as the furanocoumarin-producing clade. The lack of *5OHXS* genes in both *Morus* and *Artocarpus* supports the hypothesis that *Ficus* furanocoumarin-biosynthetic genes emerged after the *Ficus*-*Morus* divergence (Villard *et al*., 2021). The presence of orthologous *5OHXS* genes in *Ficus* and *Antiaris* indicates the 5OHXS activity emerged before the Ficeae-Olmedieae divergence. This strongly suggests that furanocoumarins were absent in the most recent *Ficus*-*Morus* ancestor but produced by the Ficeae-Olmedieae one. Based on these patterns, it is reasonable to infer that furanocoumarins emerged only once in Moraceae, during the early evolution of the furanocoumarin-producing clade. However, as metabolic and genetic data is still scarce in this plant family, we cannot yet rule out the possibility of a multiple and independent emergence of furanocoumarins in the Dorstenieae tribe and in the Ficeae-Olmedieae subclade, respectively. Additional metabolomic and genetic resources will thus be necessary to completely elucidate furanocoumarin emergence in Moraceae.

## Acknowledgements

The authors warmly thank Jessica Amaral (Universidade Federal de Sao Carlos, Brasil), Sakihito Kitajima, (Kyoto Institute of Technology, Japan), Jérémy Grosjean and Marwa Roumani (Université de Lorraine, France) for their precious help on the project, including supply in pyranocoumarins standards, access to *Ficus carica* transcriptomic data, and support in phytochemical analysis and molecular biology. The authors also address special thanks to Éléanore Lacoste, Anne-Flore Didelot, Coralie Deschamps, Yann Moreno, Hugo Schitterer, Juliette Braganti-Coral, Camille Chassin, Jeanne Couderc and Bleuenn Suard students of the Biotechnology specialization of ENSAIA, for their investment in the phylogenetic studies, CYP71B129-131 cloning, and their inputs on P450 expression. Plants were grown on the Plant Experimental Platform in Lorraine (PEPLor, Université de Lorraine, France), metabolomic analyses were conducted on the Metabolomic and Structural Analytic Platform (PASM, Université de Lorraine, France). AB was financially supported by a PhD grant provided by the French Governement and Grand Est Region. AB benefited from the mobility program DREAM provided by “Lorraine Université d’Excellence”, funded by the ANR “Investissements d’avenir” [grant number 15-004].

## Author contributions

AB, AO and CC realized the biochemical experiments. CV identified the coding sequences. CV and DRN performed phylogenetic analyses. AB and RK realized the computational structural biology. AB, CV, RL and AH wrote the article. JT, RL and AH supervised the project.

## Data availability

Nucleotide sequences coding for CYP71B129, CYP71B130 and CYP71B131a are openly available on EnsemblPlants under accession numbers FCD_00002714, FCD_00002719 and FCD_00002721 from the genome assembly UNIPI_FiCari_1.0.

