## Supplemental Files for "Using lineage-specific patterns to understand convergence of enzymatic functions led to the identification of Moraceae-specific P450s involved in furanocoumarin biosynthesis"

Article acceptance date: Click here to enter a date.

The following Supporting Information is available for this article:

**Fig. S1** Nucleotide dataset of *F. carica* and *M. notabilis* CYP71 clan sequences.

**Fig. S2** Protein sequence alignment of *F. carica* and *M. notabilis* CYP71 clan sequences.

**Fig. S3** Evaluation of the expression of CYP71Bs and associated mutants by immunodetection.

**Fig. S4** Nucleotide sequence alignment of CYP71Bs from the Nitrogen Fixing Clade.

**Fig. S5** Phylogeny of the CYP71B tribe in the Nitrogen Fixing Clade.

**Fig. S6** Restricted nucleotide sequence alignment of CYP71Bs from the Moraceae family.

**Fig. S7** Protein sequence comparison of characterized 5-hydroxyxanthotoxin synthases (5OHXSs) and closely related enzymes.

**Fig. S8** Formal identification of the xanthotoxin metabolization product by CYP71B131a_G373A and CYP71B131a_G373S.

**Fig. S9** Formal identification of the umbelliferone metabolization product by CYP71B mutants.

**Fig. S10** Determination of the optimal temperature of 5-hydroxyxanthotoxin synthases.

**Fig. S11** Determination of the optimal pH of 5-hydroxyxanthotoxin synthases.

**Fig. S12** Affinity of various CYP71Bs for xanthotoxin and umbelliferone.

**Table S1** Substrates tested for CYP71B129-131a functional screening.

**Table S2** Comparison of the *Ficus carica* and *Morus notabilis* CYPomes.

**Table S3** CYP71Bs from the Nitrogen Fixing Clade

**Fig. S1 Nucleotide dataset of *F. carica* and *M. notabilis* CYP71 clan sequences.**

(Attached Fasta dataset)

**Fig. S2 Protein sequence alignment of *F. carica* and *M. notabilis* CYP71 clan sequences**

(Attached Fasta Alignment)

**Fig. S3 Evaluation of the expression of CYP71Bs and associated mutants by immunodetection.** Western Blot performed on the microsomes collected after the expression of CYP71B129, CYP71B130, CYP71B131a, CYP71B131a_G373A, CYP71B131a_G373V, CYP71B131a_G373S and CYP71B129_A373G. Primary antibodies were directed against the His-tag added in the sequence of every candidate. The negative control (Ø plasmid) refers to the microsomes from the yeasts transformed with the empty pYeDP60_GW vector. A molecular weight marker (ProSieveTM QuadColorTM protein marker, 4.6 kDa - 300 kDa, Lonza Rockland Inc) was included.

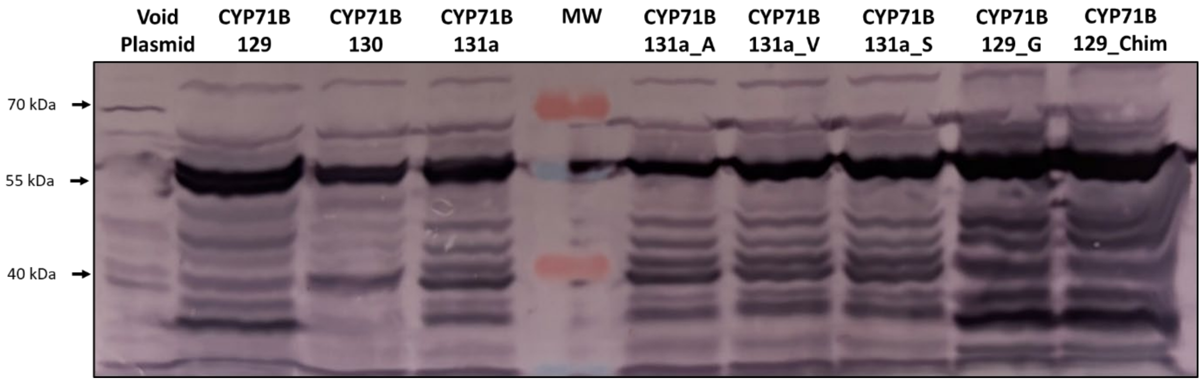


**Fig. S4 Nucleotide sequence alignment of CYP71Bs from the Nitrogen Fixing Clade.**

(Attached Fasta Alignment)

**Fig. S5 Phylogeny of the CYP71B tribe in the Nitrogen Fixing Clade.** Gene-family tree of the CYP71Bs from the Rosales (Moraceae, Urticaceae, Cannabaceae, Rhamnaeae, Rosaceae), Cucurbitales (Cucurbitaceae) and Fagales (Fagaceae), rooted on *F. carica* CYP71AH65. Sequences from the Rosales order are colored by family. Characterized 5-hydroxyxanthotoxin synthases (5OHXSs) are highlighted with a red diamond, characterized umbelliferone hydroxylase (UMBH) with a black diamond. Other characterized CYP71Bs from *P. trichocarpa*, *E. camphora*, and *A. thaliana* are described in Irmisch et al., 2014; Hansen et al., 2018 and Schuhegger et al., 2006. Remarkably, CYP71Bs from *E. camphora* and *A. thaliana* cluster with some CYP71ASs, demonstrating how the CYP71B and CYP71AS subfamilies overlap in a single clade, called the CYP71B tribe. This is consistent with the fact that, in the screened species from the Cucurbitales (*Cucumis sativus*) and Fabales (*Cajanus Cajan*, *Medicago truncatula*, not shown), no sequence was found that was closer to *F. carica* CYP71B131b than to *F. carica* CYP71AS33.

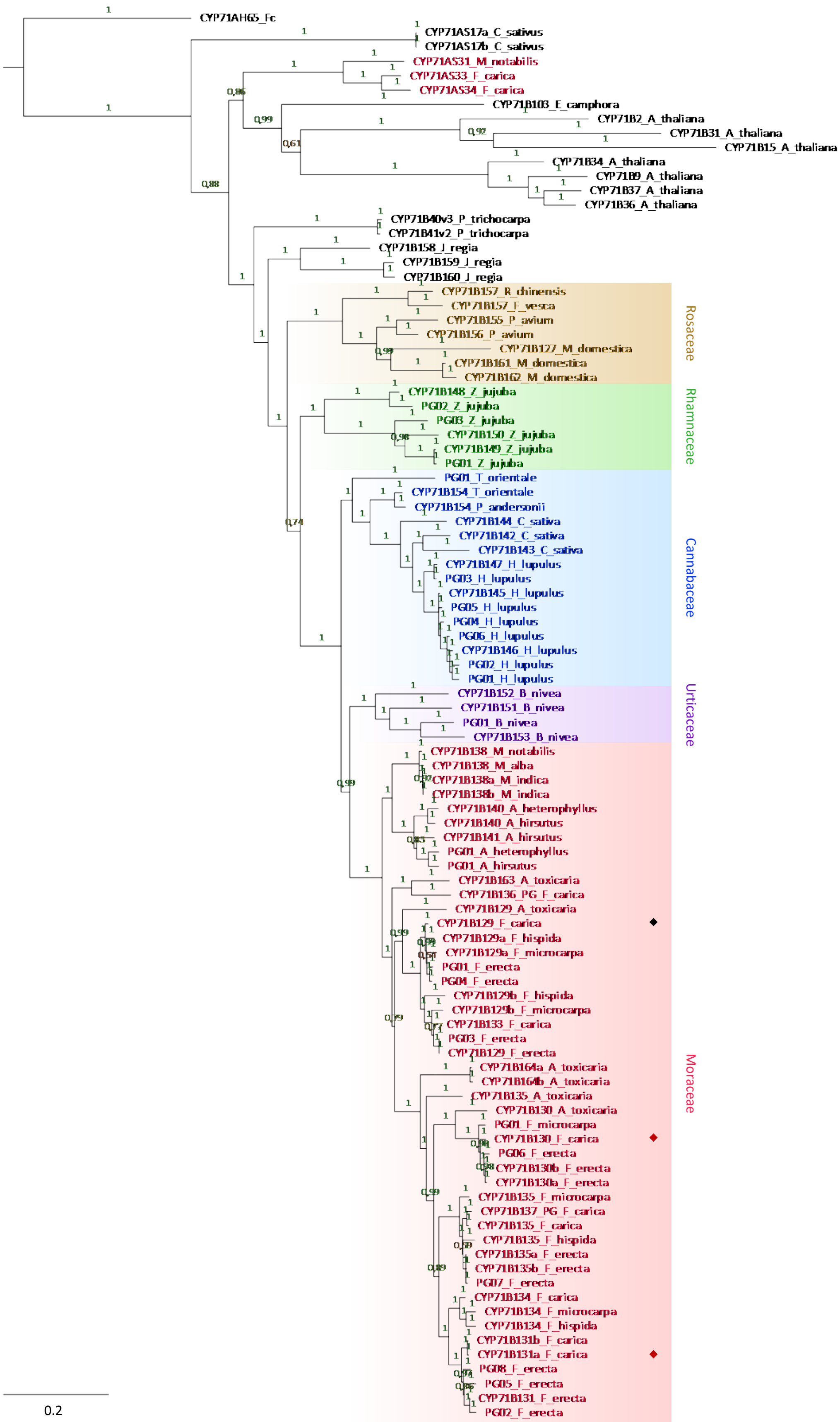
CYP71AH65_Fc
CYP71AS17a_C_sativus
CYP71AS17b_C_sativus
CYP71AS31_M_notabilis
CYP71AS33_F_carica
CYP71AS34_F_carica
CYP71B103_E_camphora
CYP71B2_A_thaliana
CYP71B31_A_thaliana
CYP71B15_A_thaliana
CYP71B34_A_thaliana
CYP71B9_A_thaliana
CYP71B37_A_thaliana
CYP71B36_A_thaliana
CYP71B40v3_P_trichocarpa
CYP71B41v2_P_trichocarpa
CYP71B158_J_regia
CYP71B159_J_regia
CYP71B160_J_regia
CYP71B157_R_chinensis
CYP71B157_F_vesca
CYP71B155_P_avium
CYP71B156_P_avium
CYP71B127_M_domestica
CYP71B161_M_domestica
CYP71B162_M_domestica
Rosaceae
CYP71B148_Z_jujuba
PG02_Z_jujuba
PG03_Z_jujuba
CYP71B150_Z_jujuba
CYP71B149_Z_jujuba
PG01_Z_jujuba
Rhamnaceae
PG01_T_orientale
CYP71B154_T_orientale
CYP71B154_P_andersonii
CYP71B144_C_sativa
CYP71B142_C_sativa
CYP71B143_C_sativa
CYP71B147_H_lupulus
PG03_H_lupulus
CYP71B145_H_lupulus
PG05_H_lupulus
PG04_H_lupulus
PG06_H_lupulus
CYP71B146_H_lupulus
PG02_H_lupulus
PG01_H_lupulus
Cannabaceae
CYP71B152_B_nivea
CYP71B151_B_nivea
PG01_B_nivea
CYP71B153_B_nivea
Urticaceae
CYP71B138_M_notabilis
CYP71B138_M_alba
CYP71B138a_M_indica
CYP71B138b_M_indica
CYP71B140_A_heterophyllus
CYP71B140_A_hirsutus
CYP71B141_A_hirsutus
PG01_A_heterophyllus
PG01_A_hirsutus
CYP71B163_A_toxicaria
CYP71B136_PG_F_carica
CYP71B129_A_toxicaria
CYP71B129_F_carica
CYP71B129a_F_hispida
CYP71B129a_F_microcarpa
PG01_F_erecta
PG04_F_erecta
CYP71B129b_F_hispida
CYP71B129b_F_microcarpa
CYP71B133_F_carica
PG03_F_erecta
CYP71B129_F_erecta
CYP71B164a_A_toxicaria
CYP71B164b_A_toxicaria
CYP71B135_A_toxicaria
CYP71B130_A_toxicaria
PG01_F_microcarpa
CYP71B130_F_carica
PG06_F_erecta
CYP71B130b_F_erecta
CYP71B130a_F_erecta
CYP71B135_F_microcarpa
CYP71B137_PG_F_carica
CYP71B135_F_carica
CYP71B135_F_hispida
CYP71B135a_F_erecta
CYP71B135b_F_erecta
PG07_F_erecta
CYP71B134_F_carica
CYP71B134_F_microcarpa
CYP71B134_F_hispida
CYP71B131b_F_carica
CYP71B131a_F_carica
PG08_F_erecta
PG05_F_erecta
CYP71B131_F_erecta
PG02_F_erecta
Moraceae
0.2

**Fig. S6 Restricted nucleotide sequence alignment of CYP71Bs from the Moraceae family.**

(Attached Fasta Alignment)

**Fig. S7 Protein sequence comparison of characterized 5-hydroxyxanthotoxin synthases (5OHXSs) and closely related enzymes.** Amino acids identical to the CYP71B130 equivalent are in blue, others are in yellow, gaps are in grey. Substrate Recognition Sites (SRSs) are highlighted.

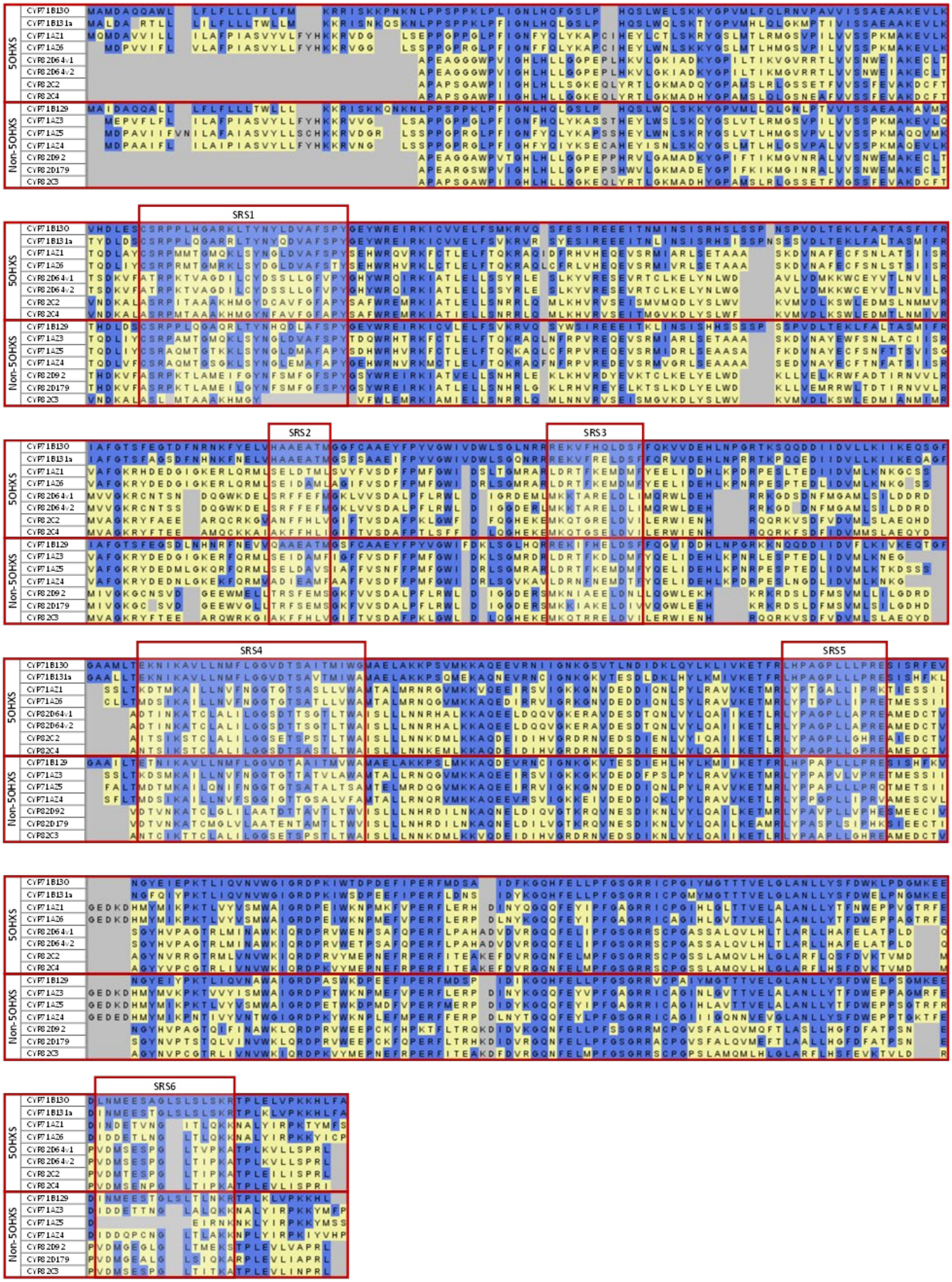

5OHXS
Non-5OHXS
CYP71B130
CYP71B131a
CYP71A21
CYP71A26
CYP82D64v1
CYP82D64v2
CYP82C2
CYP82C4
CYP71B129
CYP71A23
CYP71A25
CYP71A24
CYP82D92
CYP82D179
CYP82C3
SRS1
SRS2
SRS3
SRS4
SRS5
SRS6

**Fig. S8 Formal identification of the xanthotoxin metabolization product by CYP71B131a_G373A and CYP71B131a_G373S.** Tandem mass spectrometry fragmentation pattern of a standard of 5-hydroxyxanthotoxin (a), and of the xanthotoxin metabolization product of CYP71B131a_G373A (b) and CYP71B131a_G373S (c). Analysis was performed after negative electrospray ionisation (ESI) and Targeted Single Ion Monitoring (SIM) mode between 53 and 242 m/z. The Collision-induced Dissociation (CID) and Higher Energy Collisional Dissociation (HCD) fragmentation modes were moth investigated.

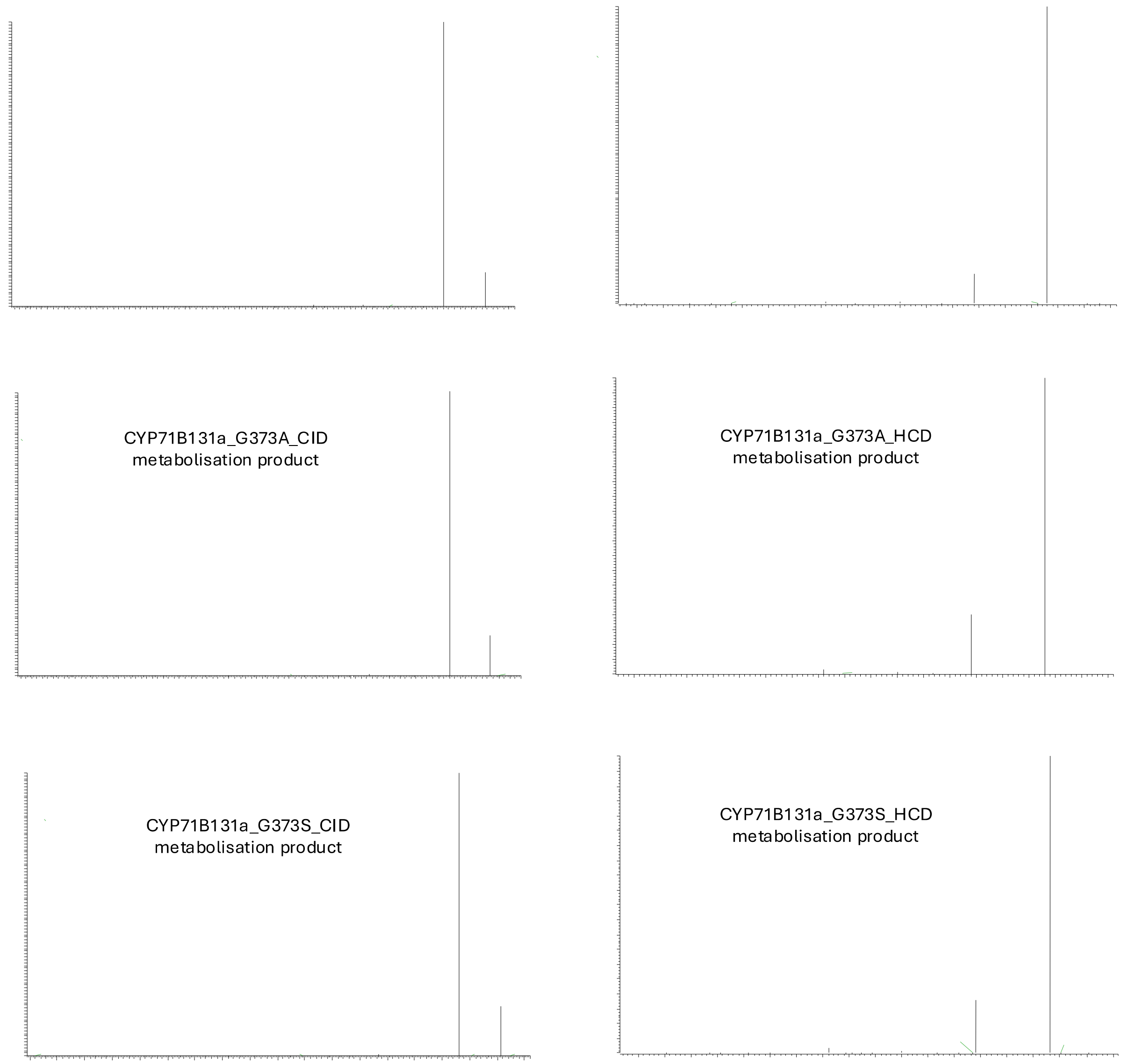

**Fig. S9 Formal identification of the umbelliferone metabolization product by CYP71B mutants.** Tandem mass spectrometry fragmentation pattern of standard of daphnetin (a), 5,7-dihydroxycoumarin (b), 4,7-dihydroxycoumarin (c) and esculetin (d), and of the umbelliferone metabolization product of CYP71B131a_G373A (e), CYP71B131a_G373S (f), CYP71B131a_G373V (g), CYP71B129_A373G (h). Analysis was performed after negative electrospray ionisation (ESI) and Targeted Single Ion Monitoring (SIM) mode between 53 and 242 m/z. The Collision-induced Dissociation (CID) and Higher Energy Collisional Dissociation (HCD) fragmentation modes were moth investigated.

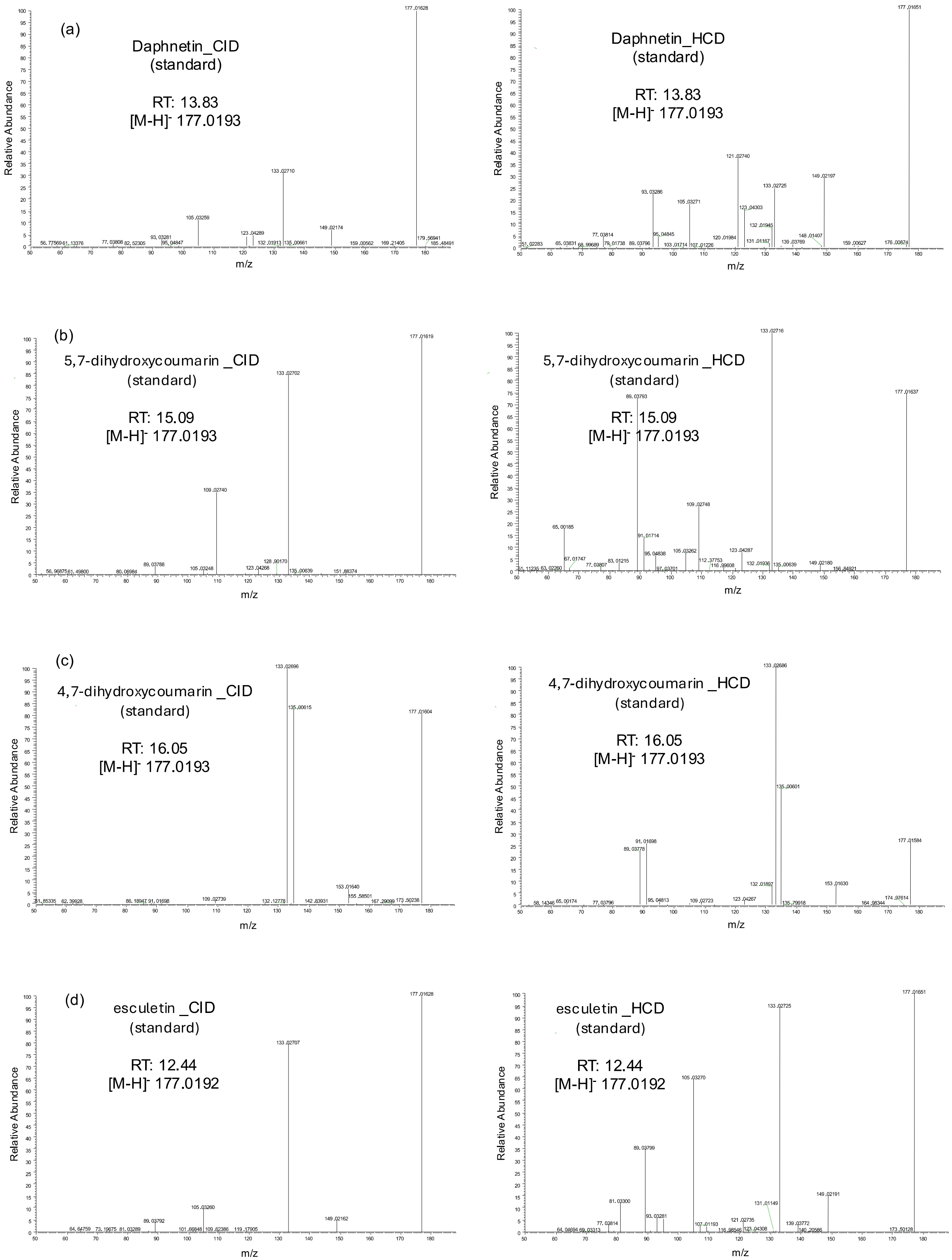
(a)
Daphnetin_CID
(standard)
RT: 13.83
[M-H]- 177.0193
Daphnetin_HCD
(standard)
RT: 13.83
[M-H]- 177.0193
(b)
5,7-dihydroxycoumarin _CID
(standard)
RT: 15.09
[M-H]- 177.0193
5,7-dihydroxycoumarin _HCD
(standard)
RT: 15.09
[M-H]- 177.0193
(c)
4,7-dihydroxycoumarin _CID
(standard)
RT: 16.05
[M-H]- 177.0193
4,7-dihydroxycoumarin _HCD
(standard)
RT: 16.05
[M-H]- 177.0193
(d)
esculetin _CID
(standard)
RT: 12.44
[M-H]- 177.0192
esculetin _HCD
(standard)
RT: 12.44
[M-H]- 177.0192
Relative Abundance
m/z

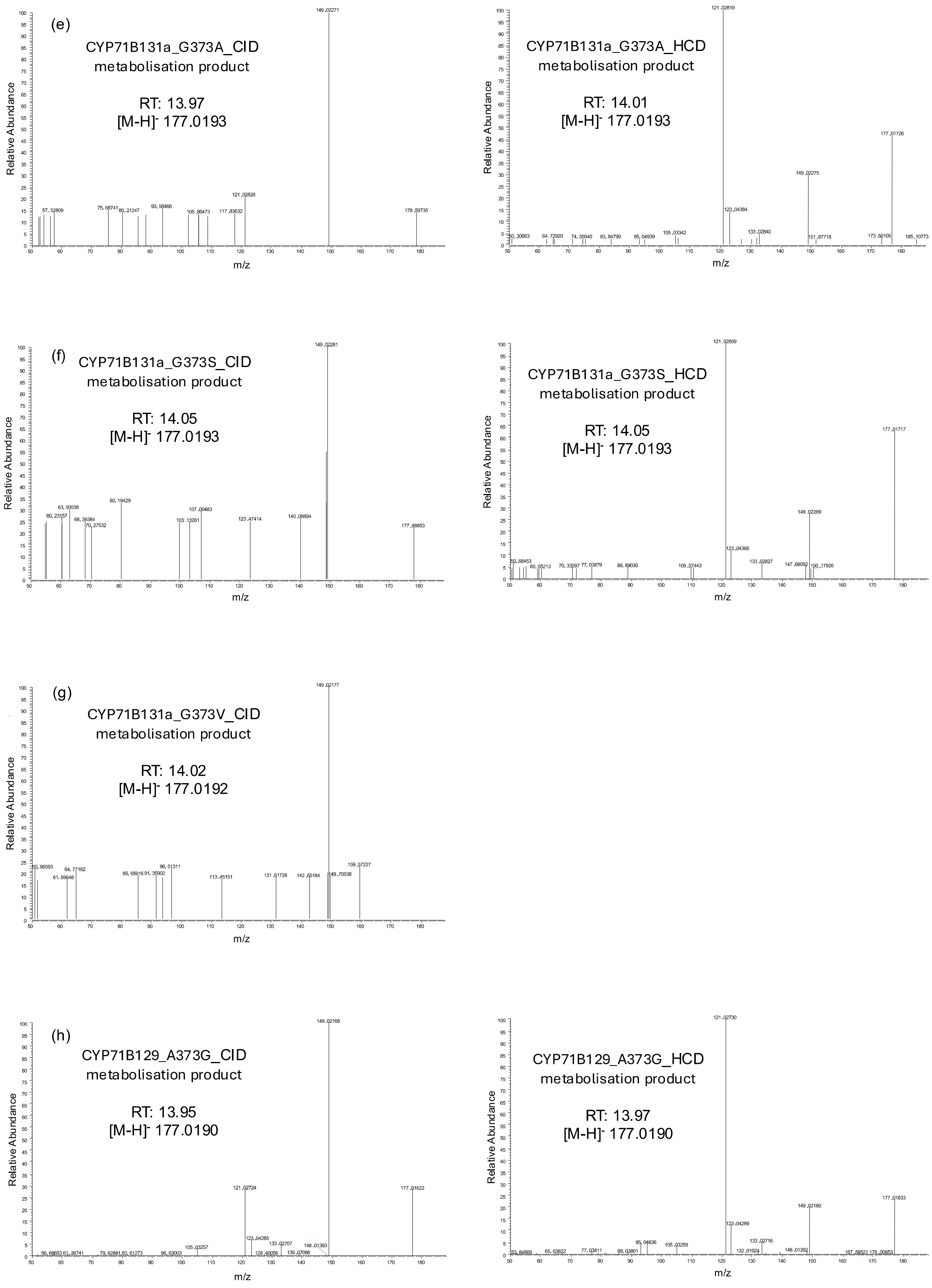

(e)
CYP71B131a_G373A_CID
metabolisation product
RT: 13.97
[M-H]- 177.0193
CYP71B131a_G373A_HCD
metabolisation product
RT: 14.01
[M-H]- 177.0193
(f)
CYP71B131a_G373S_CID
metabolisation product
RT: 14.05
[M-H]- 177.0193
CYP71B131a_G373S_HCD
metabolisation product
RT: 14.05
[M-H]- 177.0193
(g)
CYP71B131a_G373V_CID
metabolisation product
RT: 14.02
[M-H]- 177.0192
(h)
CYP71B129_A373G_CID
metabolisation product
RT: 13.95
[M-H]- 177.0190
CYP71B129_A373G_HCD
metabolisation product
RT: 13.97
[M-H]- 177.0190
Relative Abundance
m/z

**Fig. S10 Determination of the optimal temperature of 5-hydroxyxanthotoxin synthases**. Activity of CYP71B130 (a) and CYP71B131a (b) incubated in the presence of xanthotoxin and NADPH, in buffers of variable temperature. Product quantities were measured by UHPLC-MS, at 320nm. Reactions were performed in triplicates; the error bars correspond to the standard errors. The curves were plotted on SigmaPlot to fit the experimental data.

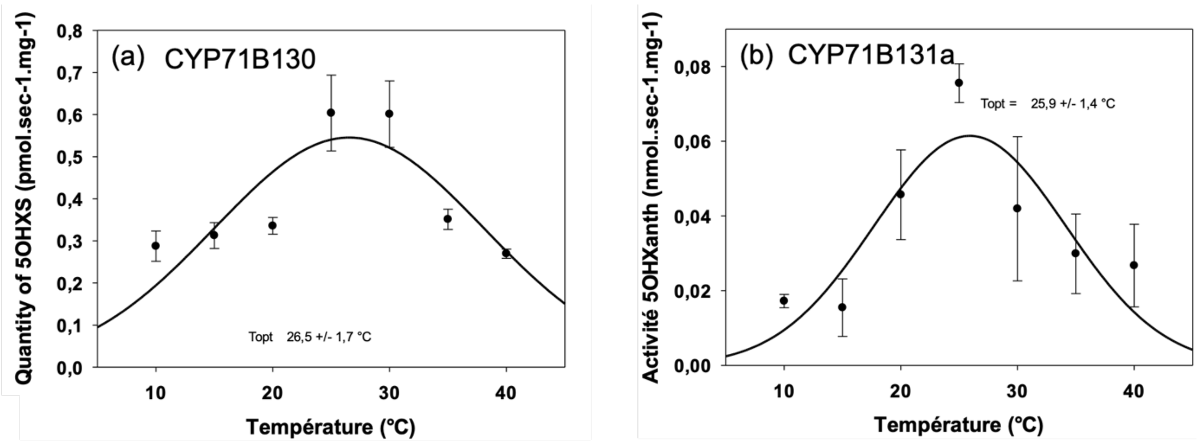


**Fig. S11 Determination of the optimal pH of 5-hydroxyxanthotoxin synthases**. Activity of CYP71B130 (a) and CYP71B131a (b) incubated in the presence of xanthotoxin and NADPH, in buffers of variable pH. Product quantities were measured by UHPLC-MS, at 320nm. Reactions were performed in triplicates; the error bars correspond to the standard errors. The curves were plotted on SigmaPlot to fit the experimental data.

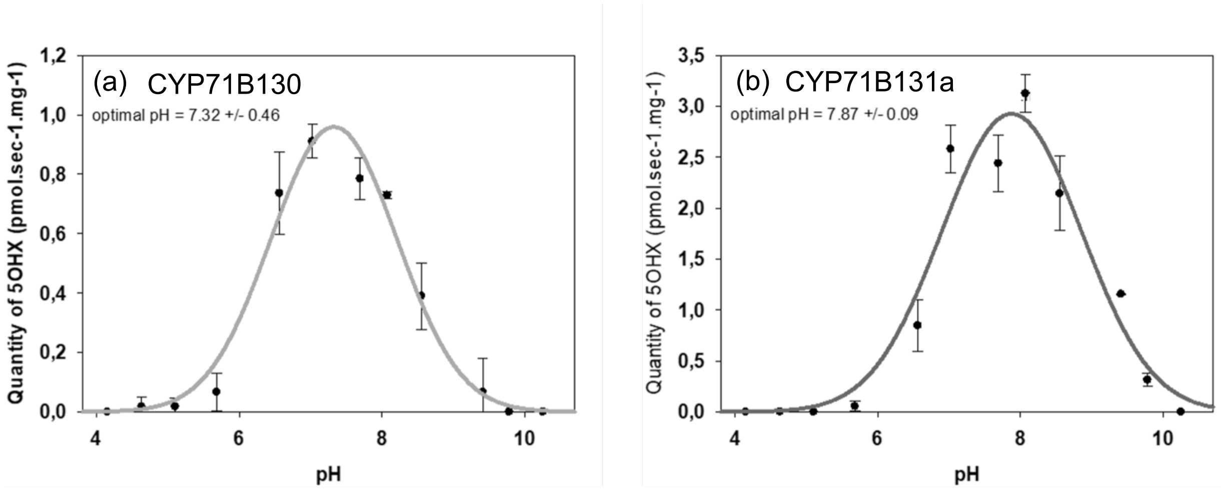

**Fig. S12 Affinity of various CYP71Bs for xanthotoxin and umbelliferone.** Specific activity of CYP71B130 (a), CYP71B131a (b) and CYP71B131a_G373A (c) in the presence of various xanthotoxin concentrations. Specific activity of CYP71B129 (d), CYP71B131a_G373A (e) and CYP71B131a_G373S (f) in the presence of various umbelliferone concentrations. Product quantities were measured by UHPLC-MS, at 320nm. Reactions were performed in triplicates; the error bars correspond to the standard errors. The Michaelis-Menten model curves were plotted on SigmaPlot to fit the experimental data. Associated kinetic parameters were determined in SigmaPlot.

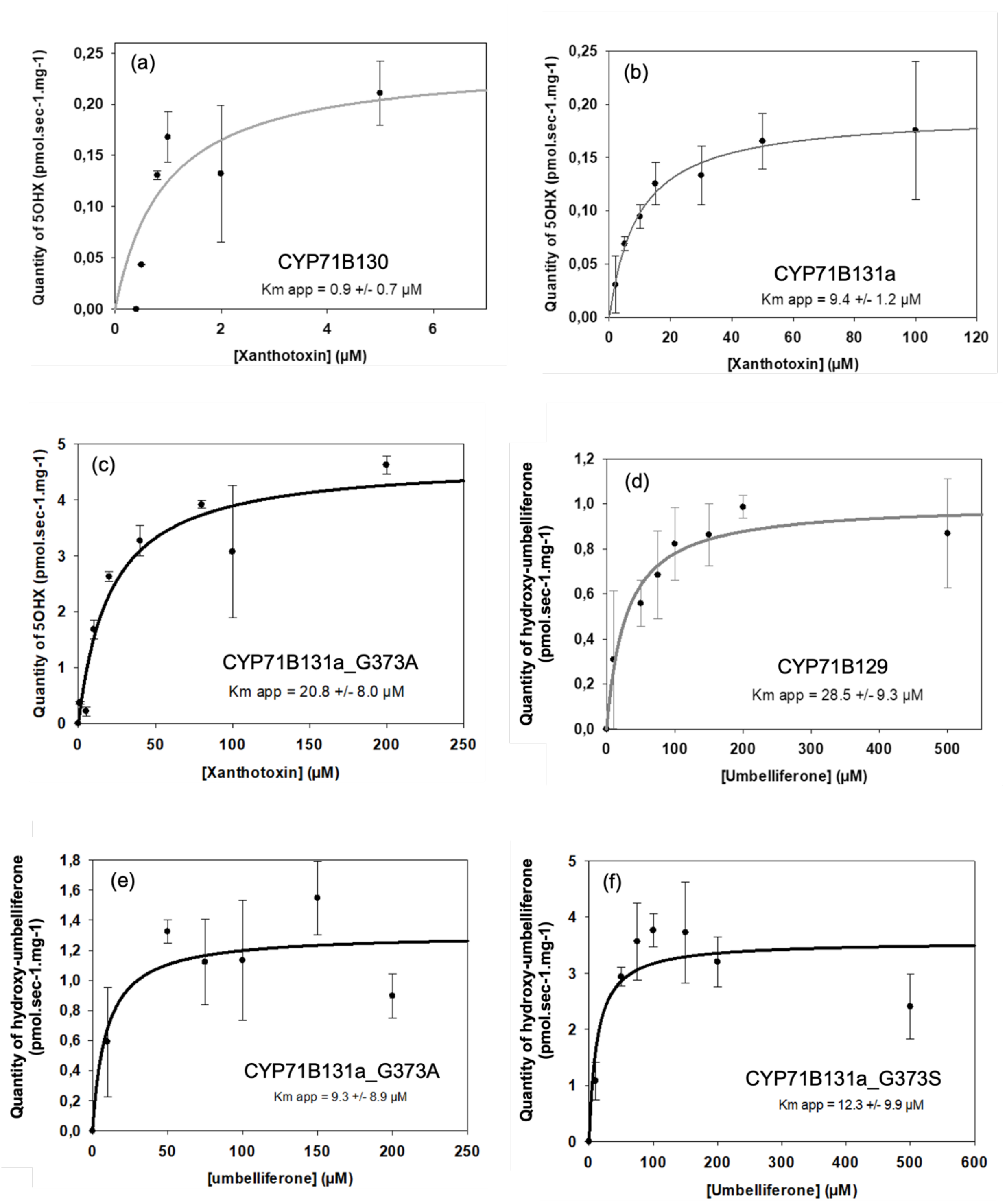


**Table S1. Substrates tested for CYP71B129-131a functional screening.**

| Family | Coumpound | Molecular Formula | Monoisotopic Mass | Provider | Reference |
|---|---|---|---|---|---|
| **Phenylpropenes** | ***t*-cinnamic acid** | C9H8O2 | 148.052429494 | Sigma | REF128708 |
| | **3,4,5-trimethoxy-cinnamic acid** | C12H14O5 | 238.08412354 | Sigma | T70408 |
| **Coumarins** | **Umbelliferone** | C9H6O3 | 162.031694049 | Extrasynthèse | REF0529 |
| | **4-methyl-umbelliferone** | C10H8O3 | 176.047344113 | Sigma | M1381 |
| | **Esculetin** | C9H6O4 | 178.02660867 | Extrasynthèse | REF0502 |
| | **Esculin** | C15H16O9 | 340.07943208 | Sigma | E8250 |
| | **3,5-dimethoxycoumarin** | C11H10O4 | 206.05790880 | Herboreal Ltd | [redacted] |
| | **5,7-dihydroxycoumarin** | C9H6O4 | 178.02660867 | Alpha chemistry | ACM2732185 |
| | **Daphnetin** | C9H6O4 | 178.02660867 | Sigma | PHL89621 |
| | **Hydrangetin** | C10H8O4 | 192.04225873 | Biosynth/Carbosynth | XD163676 |
| | **Daphnetin dimethyl-ether** | C11H10O4 | 206.05790880 | Biosynth/Carbosynth | XD164077 |
| | **Isoscopoletin** | C10H8O4 | 192.04225873 | Extrasynthèse | REF0520 |
| | **Scopoletin** | C10H8O4 | 192.04225873 | Sigma | S2500 |
| | **Scoparone** | C11H10O4 | 206.05790880 | Phytolab | REF89787 |
| | **Herniarin** | C10H8O3 | 176.047344113 | Extrasynthèse | REF0556S |
| | **4-hydroxy-herniarin** | C10H8O4 | 192.04225873 | BLD Pharmatech | MFCD00673700 |

| | | | | | |
|---|---|---|---|---|---|
| | 6,7,8-trihydroxycoumarin | C9H6O5 | 194.02152329 | Green pharma | REF5718418 |
| | Fraxetin | C10H8O5 | 208.03717335 | Phytolab | REF89549 |
| | Fraxin | C16H18O10 | 370.08999677 | Phytolab | REF89545 |
| | Dimethyl-fraxetin | C12H12O5 | 236.06847348 | MedChemExpress | HY-N0085 |
| | Fraxidin | C11H10O5 | 222.05282342 | Extrasynthèse | REF0525 |
| | Isofraxidin | C11H10O5 | 222.05282342 | Biosynth/Carbosynth | FI65562 |
| | Suberosin | C15H16O3 | 244.109944368 | Herboreal Ltd | [redacted] |
| | Demethylsuberosin | C14H14O3 | 230.094294304 | R. Munakata | |
| | Osthol | C15H16O3 | 244.109944368 | Extrasynthèse | REF0541S |
| | o-prenyl-umbelliferone | C14H14O3 | 230.094294304 | Biosynth/Carbosynth | KAA38750 |
| | Aurapten | C19H22O3 | 298.15689456 | Extrasynthèse | REF0544 |
| | 5-geranyloxy-7-methoxycoumarin | C20H24O4 | 328.16745924 | Extrasynthèse | REF0560S |
| | 6-methoxycoumarin | C10H8O3 | 176.047344113 | Combi-Blocks | HA-2519 |
| | Coumarin | C9H6O2 | 146.036779430 | Sigma | C4261 |
| | 8-methoxycoumarin | C10H8O3 | 176.047344113 | CymitQuimica | TR-M299275 |
| | Limettin | C11H10O4 | 206.05790880 | Sigma | REF116238 |

| | | | | | |
|---|---|---|---|---|---|
| | 5-methoxy-7-hydroxycoumarin | C10H8O4 | 192.04225873 | MedChemExpress | HY-N7179 |
| | 5-hydroxy-7-methoxycoumarin | C10H8O4 | 192.04225873 | Biosynth/Carbosynth | YAA05361 |
| | 6,7,8-trihydroxycoumarin | C9H6O5 | 194.02152329 | Green pharma | REF5718418 |
| Linear furanocoumarins | Psoralen | C11H6O3 | 186.031694049 | Extrasynthèse | REF0553S |
| | Xanthotoxol | C11H6O4 | 202.02660867 | Extrasynthèse | REF0534 |
| | Bergaptol | C11H6O4 | 202.02660867 | Extrasynthèse | REF0590S |
| | Xanthotoxin | C12H8O4 | 216.04225873 | Sigma | M3501 |
| | Bergapten | C12H8O4 | 216.04225873 | Extrasynthèse | REF503 |
| | 5-Hydroxyxanthotoxin | C12H8O5 | 232.03717335 | Herboreal Ltd | [redacted] |
| | 8-hydroxybergapten | C12H8O5 | 232.03717335 | Herboreal Ltd | [redacted] |
| | Isopimpinellin | C13H10O5 | 246.05282342 | Extrasynthèse | REF0536S |
| | Marmesin | C14H14O4 | 246.08920892 | TransMIT | M034 |
| | Celereoin | C14H14O5 | 262.08412354 | Prof. W. Boland (Jena, Germany) | [redacted] |
| | Imperatorin | C16H14O4 | 270.08920892 | Herboreal Ltd | [redacted] |
| | Isoimperatorin | 16H14O4 | 270.08920892 | Herboreal Ltd | [redacted] |
| | Heraclenin | C16H14O5 | 286.08412354 | Herboreal Ltd | [redacted] |
| | Heraclenol | C16H16O6 | 304.09468823 | Herboreal Ltd | [redacted] |

| | | | | | |
|---|---|---|---|---|---|
| | **Byakangelicol** | $C_{17}H_{16}O_6$ | 316.09468823 | Herboreal Ltd | |
| | **Byakangelicin** | $C_{17}H_{18}O_7$ | 334.10525291 | Herboreal Ltd | |
| | **8-Geranyloxypsoralen** | $C_{21}H_{22}O_4$ | 338.15180918 | Ryosuke Munakata-KYOTO | |
| | **Bergamottin** | $C_{21}H_{22}O_4$ | 338.15180918 | Extrasynthèse | REF0550S |
| | **Epoxybergamottin** | $C_{21}H_{22}O_5$ | 354.14672380 | Herboreal Ltd | |
| | **6,7-dihydroxybergamottin** | $C_{21}H_{24}O_6$ | 372.15728848 | Herboreal Ltd | |
| | **Cnidilin** | $C_{17}H_{16}O_5$ | 300.09977361 | Herboreal Ltd | |
| | **Cnidicin** | $C_{21}H_{22}O_5$ | 354.14672380 | HerborealLtd | |
| | **Swietenocoumarin B** | $C_{17}H_{16}O_4$ | 284.10485899 | AnalytiCon Discovery, GmbH | NP-013575 |
| | **Chalepensin** | $C_{16}H_{14}O_3$ | 254.094294304 | AnalytiCon Discovery, GmbH | NP-002897 |
| | **Phellopterin** | $C_{17}H_{16}O_5$ | 300.09977361 | Herboreal Ltd | |
| | **Rutarin** | $C_{20}H_{24}O_{10}$ | 424.13694696 | AnalytiCon Discovery, GmbH | NP-002712 |
| | **Oxypeucedanin** | $C_{16}H_{14}O_5$ | 286.08412354 | Herboreal Ltd | |
| | **Oxypeucedanin hydrate** | $C_{16}H_{16}O_6$ | 304.09468823 | Herboreal Ltd | |
| **Angular furanocoumarins** | **Angelicin** | $C_{11}H_6O_3$ | 186.031694049 | Extrasynthèse | REF0537S |
| | **Sphondinol** | $C_{11}H_6O_4$ | 202.02660867 | Herboreal | |

| | | | | | |
|---|---|---|---|---|---|
| | **Isobergapten** | C12H8O4 | 216.04225873 | Herboreal Ltd | |
| | **6-Isopentenyloxyisobergapten** | C17H16O5 | 300.09977361 | Herboreal Ltd | |
| | **Sphondin** | C12H8O4 | 216.04225873 | Herboreal Ltd | |
| | **Pimpinellin** | C13H10O5 | 246.05282342 | Extrasynthèse | REF0536S |
| | **Columbianetin** | C14H14O4 | 246.08920892 | TransMIT | C078 |
| | **Columbianadin** | C19H20O5 | 328.13107373 | Sigma | PHL83278 |
| **Pyranocoumarins** | **Xanthyletin** | C14H12O3 | 228.078644241 | Jessica Amaral-BRAZIL | |
| | **Seselin** | C14H12O3 | 228.078644241 | Jessica Amaral-BRAZIL | |
| **Others** | **7-methoxybenzofuran** | C9H8O2 | 148.052429494 | CymitQuimica | AN-AG0035YX-1g |
| | **5,6-Dihydro-2H-pyran-2-one** | C5H6O2 | 98.036779430 | CymitQuimica | 3B-D2261 |
| | **Furan** | C4H4O | 68.026214747 | CymitQuimica | 02-A13102 |

**Table S2** Comparison of the Ficus carica and Morus notabilis CYPomes. (a) List and characteristics of the CYP71 clan sequences identified in the F. carica genome. (b) Summary: classification of the F. carica and M. notabilis CYP71 clan sequences.

(Attached Excel File)

**Table S3** CYP71Bs from the Nitrogen Fixing Clade. (a) List and characteristics of the CYP71Bs. Sequences were coloured according to the classification of the associated plant species. Species from the Moraceae family (Rosales order) are in green. Species from another family of the Rosales order are in yellow. Species from another order of the Nitrogen Fixing Clade are in orange. When only genomic data were available, introns were predicted and manually removed (m.r). (b) Substrate recognitions sites (SRSs) of CYP71B129-131a. Residues are numbered according to the CYP71B131a equivalent.

(Attached Excel File)
